# Complexin cooperates with Bruchpilot to tether synaptic vesicles to the active zone cytomatrix

**DOI:** 10.1101/350207

**Authors:** Nicole Scholz, Nadine Ehmann, Divya Sachidanandan, Cordelia Imig, Benjamin H. Cooper, Olaf Jahn, Kerstin Reim, Nils Brose, Martin Pauli, Manfred Heckmann, Christian Stigloher, Tobias Langenhan, Robert J. Kittel

## Abstract

Information processing by the nervous system depends on the release of neurotransmitter from synaptic vesicles (SVs) at the presynaptic active zone. Molecular components of the cytomatrix at the active zone (CAZ) regulate the final stages of the SV cycle preceding exocytosis and thereby shape the efficacy and plasticity of synaptic transmission. Part of this regulation is reflected by a physical association of SVs with filamentous CAZ structures. However, our understanding of the protein interactions underlying SV tethering by the CAZ is far from complete. The very C-terminal region of Bruchpilot (Brp), a key component of the *Drosophila* CAZ, participates in SV tethering. Yet so far, no vesicular or cytoplasmic molecules have been reported to engage in an interaction with Brp’s C-terminus. Here, we carried out an in vivo screen for molecules that link the Brp C-terminus to SVs. This strategy identified the conserved SNARE (soluble *N*-ethylmaleimide-sensitive factor (NSF) attachment protein receptor) regulator Complexin (Cpx) as a vesicular interaction partner of Brp. We show that Brp and Cpx interact genetically and functionally. Interfering with Cpx targeting to SVs mirrored distinctive features of a C-terminal Brp truncation: impaired SV recruitment to the CAZ and enhanced short-term synaptic depression. Extending the study beyond *Drosophila* synapses, we interrogated active zones of mouse rod bipolar cells. Here, too, we collected evidence for an evolutionarily conserved role of Cpx upstream of SNARE complex assembly where it participates in SV tethering to the CAZ.

## INTRODUCTION

In the nervous system, information is represented and processed by means of neuronal action potentials (APs). The ability of neurons to fire APs at high frequency places challenging demands on chemical synapses. To sustain the speed and temporal precision of synaptic transmission, presynaptic terminals must rapidly reload SVs at the active zone and prime them for exocytosis. During high frequency stimulation, synapses often display short-term depression due to a transient drop in presynaptic neurotransmitter release. Many aspects of this phenomenon can be described by a limited pool of readily-releasable vesicles (RRVs) at the active zone membrane, which is rapidly exhausted and then refilled from larger supply pools (Hallermann et al., 2010a; Neher, 2015; Zucker and Regehr, 2002). The protein-rich CAZ appears to play an important role in regulating such short-term synaptic plasticity by guiding SV replenishment (Fernández-Busnadiego et al., 2013; Hallermann and Silver, 2013; Midorikawa and Sakaba, 2015). However, very little is known about the molecular mechanisms of SV reloading and the protein interactions that link SVs to the CAZ. This is because functional recordings of exo- and endocytosis provide only indirect information on processes preceding transmitter release and low affinity, transient interactions between SVs and the CAZ, which may be required for rapid vesicle fusion, can easily escape biochemical detection.

Brp is an essential protein component of the *Drosophila* CAZ (Kittel et al., 2006; Wagh et al., 2006). It shapes the filamentous CAZ structure by assembling as long polarized oligomers with its N-terminus near Ca^2+^-channels at the active zone membrane and its C-terminus extending into the cytoplasm (Ehmann et al., 2014; Fouquet et al., 2009). Functionally, Brp-dependent CAZ assembly is required for proper Ca^2+^-channel clustering to ensure adequate neurotransmitter release probability (Kittel et al., 2006). Moreover, the very C-terminal region of Brp tethers SVs to the cytomatrix. At synapses of *brp^nude^* mutants, which lack the 17 C-terminal amino acids of Brp (~1% of the protein), disrupted SV tethering is accompanied by short-term synaptic depression, impaired sustained transmitter release, and a slowed recovery phase (Hallermann et al., 2010b). Thus, Brp helps to establish release sites and accelerates the recruitment of SVs, enabling rapid and efficient excitation-secretion coupling at the active zone.

This basic understanding of Brp function provides an entry point to study molecular mechanisms of SV tethering to the CAZ and to shed light on protein interactions, which sustain ongoing synaptic transmission. Here, we devised an in vivo screen to search for vesicular interaction partners of Brp including those with low affinity. Surprisingly, our results show that Cpx, a key regulator of the core fusion machinery, participates in the SV cycle upstream of exocytosis. Besides interacting with the assembled *trans*-SNARE complex, this small, multifunctional protein also links SVs to Brp filaments and supports rapid SV recruitment to prevent short-term synaptic depression.

## RESULTS

### Expression of Brp peptides in motoneurons alters SV localization

The 17 C-terminal amino acids of Brp (Brp^C-tip^ hereafter) are required for efficient SV tethering to the CAZ (Hallermann et al., 2010b). We therefore tested whether a peptide encoding this amino acid sequence would in turn localize to SVs. To this end, we utilized the bipartite *GAL4-UAS* expression system (Brand and Perrimon, 1993) to drive a CFP (cyan fluorescent protein) and FLAG-tagged fusion construct of Brp^C-tip^ in the cytoplasm of glutamatergic larval *Drosophila* motoneurons (**Figure 1A, B**; *ok6>3xFlag::CFP::brp^C-tip^; abbreviated brp^C-tip^*). As would be expected for a SV-bound peptide, immunostainings of the larval neuromuscular junction (NMJ) showed a clear signal overlap of Brp^C-tip^ with the *Drosophila* vesicular glutamate transporter (VGlut; **Figure 1C**) (Daniels, 2004).

**Figure 1.**
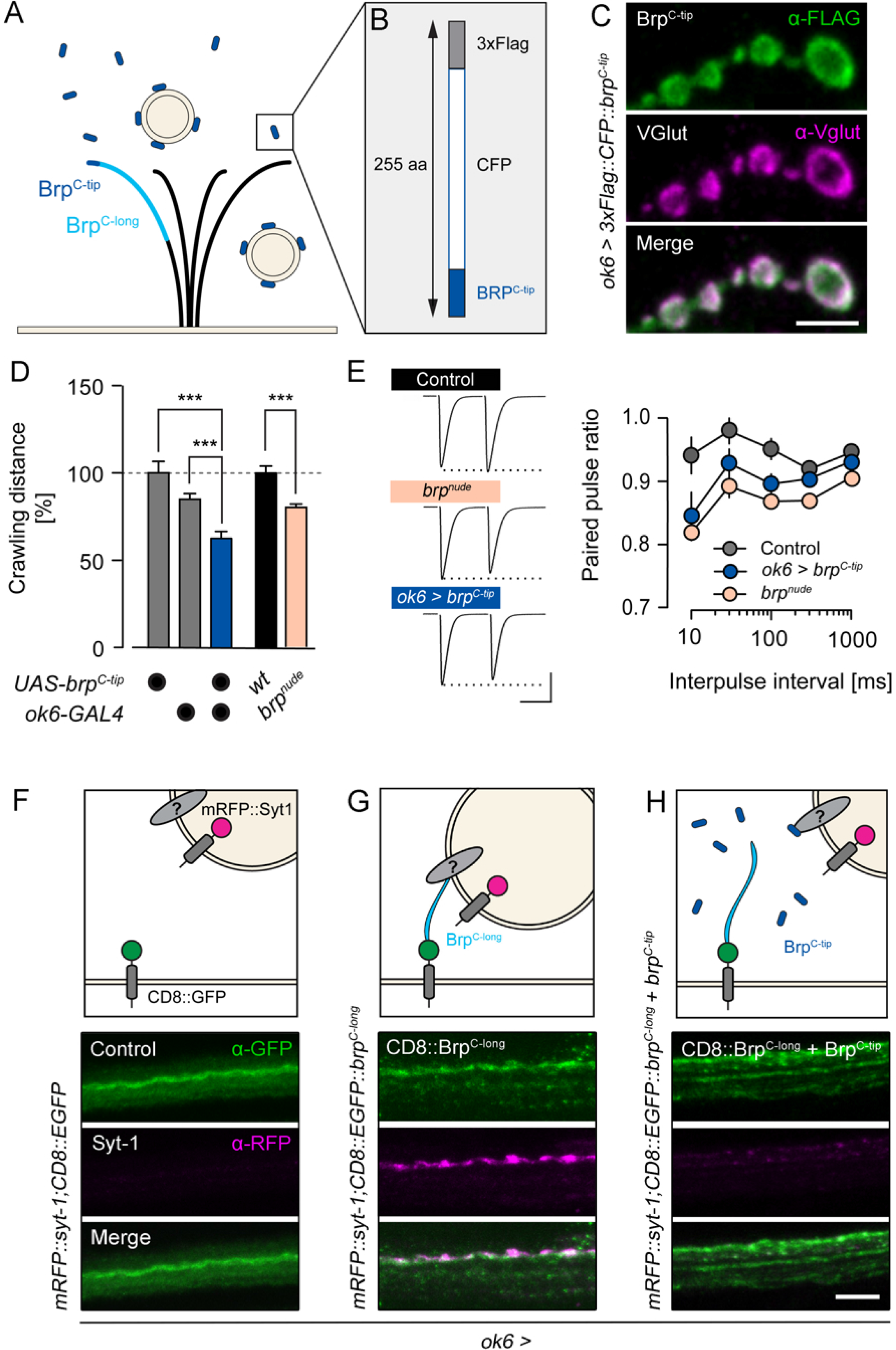
Neuronally expressed Brp peptides modify SV targeting. **(A)** Brp adopts a polarized orientation (light blue, ~ C-terminal half) to tether SVs near the AZ membrane. **(B)** A peptide containing the last 17 C-terminal amino acids of Brp (dark blue, Brp^C-tip^) fused to CFP and a FLAG-tag binds SVs. **(C)** Genetically expressed Brp^C-tip^ (green, α-FLAG, *ok6-GAL4* driver) co-localizes with SVs (magenta, α-VGlut) in the bouton cortex of motoneurons and **(D)** mimics the impaired locomotion and **(E)** paired-pulse depression of *brp^nude^* mutants. **(F-H)** Upper panels: schematic illustrations of Brp-dependent SV enrichment in the axon. Lower panels: larval motor axons co-expressing mRFP::Syt-1 with **(F)** CD8::EGFP, **(G)** CD8::EGFP::Brp^C-long^, and **(H)** CD8::EGFP::Brp^C-long^ + Brp^C-tip^. Maximal projections of confocal stacks stained against GFP (green) and RFP (magenta). Data presented as mean ± SEM, *** P ≤ 0.001. Scale bars (C) 3 μm, (E) 40 nA, 20 ms, (F-H) 5 μm.

We reasoned that if Brp interaction partners exist on SVs, then overexpressed cytoplasmic Brp^C-tip^ would compete with endogenous, active zone resident Brp for the relevant vesicular binding sites. Functionally, this should phenocopy a C-terminal Brp truncation. *brp^nude^* larvae display impaired locomotion and pronounced short-term synaptic depression (**Figure 1D, E**) (Hallermann et al., 2010b). Consistent with obstructed SV binding by Brp at the active zone, neuronal expression of Brp^C-tip^ also decreased larval locomotion (**Figure 1D**, **S1A**, **Table S1**) and enhanced paired-pulse depression of AP-evoked excitatory postsynaptic currents (eEPSCs) at short interpulse intervals (**Figure 1E, Table S2**; rank sum test versus Control: *brp^nude^*, 10 ms P = 0.0014, 30 ms P = 0.0086, all other intervals P ≤ 0.001; *brp^C-tip^*, 30 ms P = 0.0351, 100 ms P = 0.0120, all other intervals n.s.). Collectively, these results show that Brp^C-tip^ binds SVs.

Based on these findings, we investigated whether membrane-anchored transgenic Brp variants could be used to enrich SVs at ectopic sites. In order to localize Brp fragments to the axonal membrane distant from the NMJ, C-terminal segments of different length were fused to GFP-tagged CD8 (*CD8::EGFP::brp^C-x^*). For visualization, SVs were co-labelled by Synaptotagmin-1 fused to an mRFP moiety (*mRFP::syt-1*) and both genetic constructs were expressed in motoneurons. Without the addition of a C-terminal Brp fragment, membrane-associated CD8::EGFP did not lead to the accumulation of SVs in motoneuron axons (**Figure 1F**). Including the short Brp^C-tip^ sequence in the recombinant protein also failed to concentrate vesicles in the axon (**Figure S1B**). In contrast, when a longer fragment comprising roughly the C-terminal half of Brp (Brp^C-long^; **Figure 1A**) was fused to the CD8::EGFP carrier protein, SVs accumulated in the axon (**Figure 1G**). To control whether this effect is specifically mediated by membrane-anchored Brp^C-long^ we co-expressed cytoplasmic Brp^C-tip^ and assayed ectopic SV localization. As expected, axonal enrichment of SVs was abolished (**Figure 1H**). This demonstrates that cytoplasmic and membrane-bound Brp fragments containing the C-terminal 17 amino acid peptide sequence compete with each other for SV targets (**Figure 1H**).

### Forward genetic screen for a Brp interaction partner

We made use of membrane-anchored Brp^C-long^ and the concomitant SV clusters in the axon to screen for vesicular interaction partners of the Brp C-terminus through RNA interference (RNAi). This strategy comprised scoring a selection of fly strains containing *UAS-RNAi* constructs, which interfere with mRNAs encoding presynaptic and vesicle-associated gene products (**Table S3**). Knockdown of molecules that mediate an interaction between Brp and the SV should disrupt complex formation by CD8::EGFP::Brp^C-long^ and mRFP::Syt-1 at ectopic sites within the axon. This can be readily detected with light microscopy (**Figure 2A**). Importantly, screening takes place under physiological conditions, which should allow to capture protein interactions with low affinity and specific molecular requirements in vivo. The screen also facilitates the identification of indirect interactors, i.e. molecules that mediate the link between Brp and the SV without necessarily being contacted by either molecule itself. Thus, we expected our screening protocol to exceed the sensitivity of standard biochemical approaches in its ability to identify functionally relevant molecular interactions at the CAZ-SV interface.

**Figure 2.**
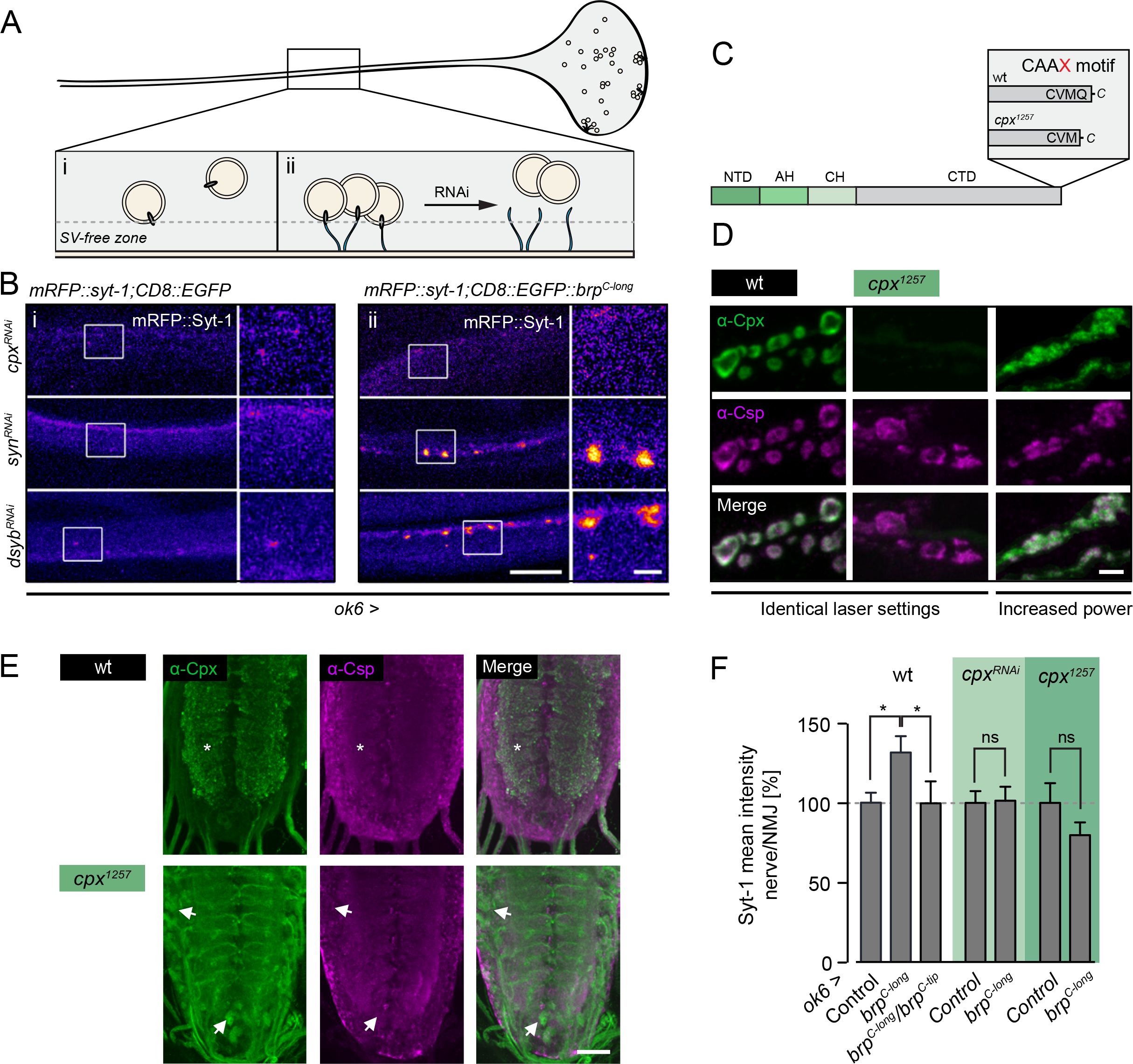
Fluorescence-based genetic screen for tethering partners of Brp. **(A)** Screening strategy. (i) The density of SVs is low in the motoneuron axon and (ii) increases upon tethering to membrane-anchored Brp (CD8::EGFP::Brp^C-long^). RNAi-mediated knockdown of factors mediating the SV-Brp interaction should prevent the axonal accumulation of SVs. **(B)** Confocal images of SVs (mRFP::Syt-1) in motoneuron axons in the absence (left, CD8::EGFP) and presence (right, CD8::EGFP::Brp^C-long^) of membrane-bound Brp (*ok6-GAL4* driver). *Cpx^RNA^* prevents Brp-dependent localization of SVs in the axon, whereas other RNAi lines, e.g. *syn^RNAi^* and *dysb^RNAi^*, have no discernible effect (**Table S3**). **(C)** Cpx layout, N-terminal domain (NTD), accessory helix (AH), central helix (CH), C-terminal domain (CTD). Deletion of the terminal glutamine disrupts the CAAX-motif in the *cpx^1257^* mutant. **(D)** Maximal projections of confocal stacks show co-localization of Cpx (green, α-Cpx) with SVs (magenta, α-Csp) in wt motoneuron boutons (left). Cpx staining intensity is strongly reduced in *cpx^1257^* boutons (centre) and no longer matches the SV distribution (right, increased laser power). **(E)** Whereas Cpx is prominent in synapse-rich regions of the wt VNC, Cpx is mainly found in somata of *cpx^1257^* mutants (arrows). **(F)** Quantification of nerve/NMJ ratio of the mRFP::Syt signal. The BRP^C-long^-dependent increase of SVs in the nerve is prevented by *brp^C-tip^* and *cpx^RNAi^* expression (*ok6-GAL4*) and in the *cpx^1257^* mutant. Data presented as mean ± SEM, * P ≤ 0.05. Scale bars (B) 10 μm, 2 μm (inset), (D) 3 μm, (E) 30 μm.

We tested 27 candidate genes (**Table S3**). Whereas knockdown of most targets did not noticeably perturb the Brp^C-long^-dependent enrichment of SVs in axons [examples shown for *dysbindin* (*dysb*) and *synapsin* (*syn*); **Figure 2B**], motoneuronal expression of an RNAi construct directed against *cpx* (*ok6>cpx^RNAl^*) reduced axonal SV levels (**Figures 2B**, **S2**). We therefore studied a possible link between Cpx, SVs, and Brp in more detail.

The C-terminus of the predominant Cpx isoform in *Drosophila* (Cpx7A) contains a membrane binding farnesylation site (**Figure 2C**, CAAX-motif; CVMQ) (Buhl et al. 2013; Zhang and Casey, 1996). This motif is important for clamping spontaneous fusion events and for localizing Cpx to sites of exocytosis, possibly through an association with SVs (Buhl et al. 2013; Iyer et al., 2013). We therefore chose a *cpx* mutant, *cpx^1257^*, which lacks the last C-terminal glutamine and exhibits disrupted farnesylation (Iyer et al., 2013) to further investigate Cpx in the context of SV interactions (**Figure 2C**). At the wild-type (wt) NMJ, Cpx is distributed evenly around the bouton cortex, similar to vesicle proteins such as SV-associated Csp (cysteine string protein; **Figures 2D**) (Buhl et al. 2013). In contrast, *cpx^1257^* mutants (*cpx^1257^/cpx^Sh1^*) displayed strongly reduced Cpx levels at the NMJ (**Figure 2D**). Here, the protein formed clusters primarily next to Brp profiles, indicating disturbed synaptic localization (Buhl et al. 2013). Cpx enrichment at the cortex was drastically decreased and the signal in inter-bouton regions and the main nerve fascicle appeared proportionally increased (**Figures 2D**, **S3**, **S4**). In the wt ventral nerve chord (VNC), Cpx and Csp signals partially overlapped. In the *cpx^1257^* mutant, however, Cpx was mainly concentrated in cell bodies (**Figure 2E**). Consistent with previous work (Buhl et al. 2013; Iyer et al., 2013), these results suggest that C-terminal farnesylation of *Drosophila* Cpx promotes its association with SVs and its targeting to presynaptic terminals.

We quantified the contribution of Cpx to axonal SV targeting by Brp^C-long^. In the wt background, the ratio of SVs located in the nerve vs. the NMJ was significantly increased upon expression of membrane-anchored Brp^C-long^ and reverted by co-expression of cytoplasmic Brp^C-tip^ (**Figure 2F**). Importantly, the nerve/NMJ ratio of SVs was levelled out in *cpx^1257^* mutants and by *cpx^RNAi^* (**Figure 2F**, **Table S4**). These results indicate that Cpx supports SV tethering to Brp.

### Cpx and Brp interact genetically

To substantiate these findings, we examined the genetic relationship between *cpx* and *brp*. Beginning with a classical epistasis assay, we tested locomotion of single (*brp^nude^/brp^69^* or *cpx^1257^/cpx^Sh1^*) and double mutant (‘DM’, *brp^nude^/brp^69^*;*cpx^1257^/cpx^Sh1^*) larvae. Both *brp^nude^* and *cpx^1257^* animals displayed crawling defects. Notably, these phenotypes were not additive, implying that both proteins act in the same pathway at the synapse (**Figure 3A**, **Table S1**).

**Figure 3.**
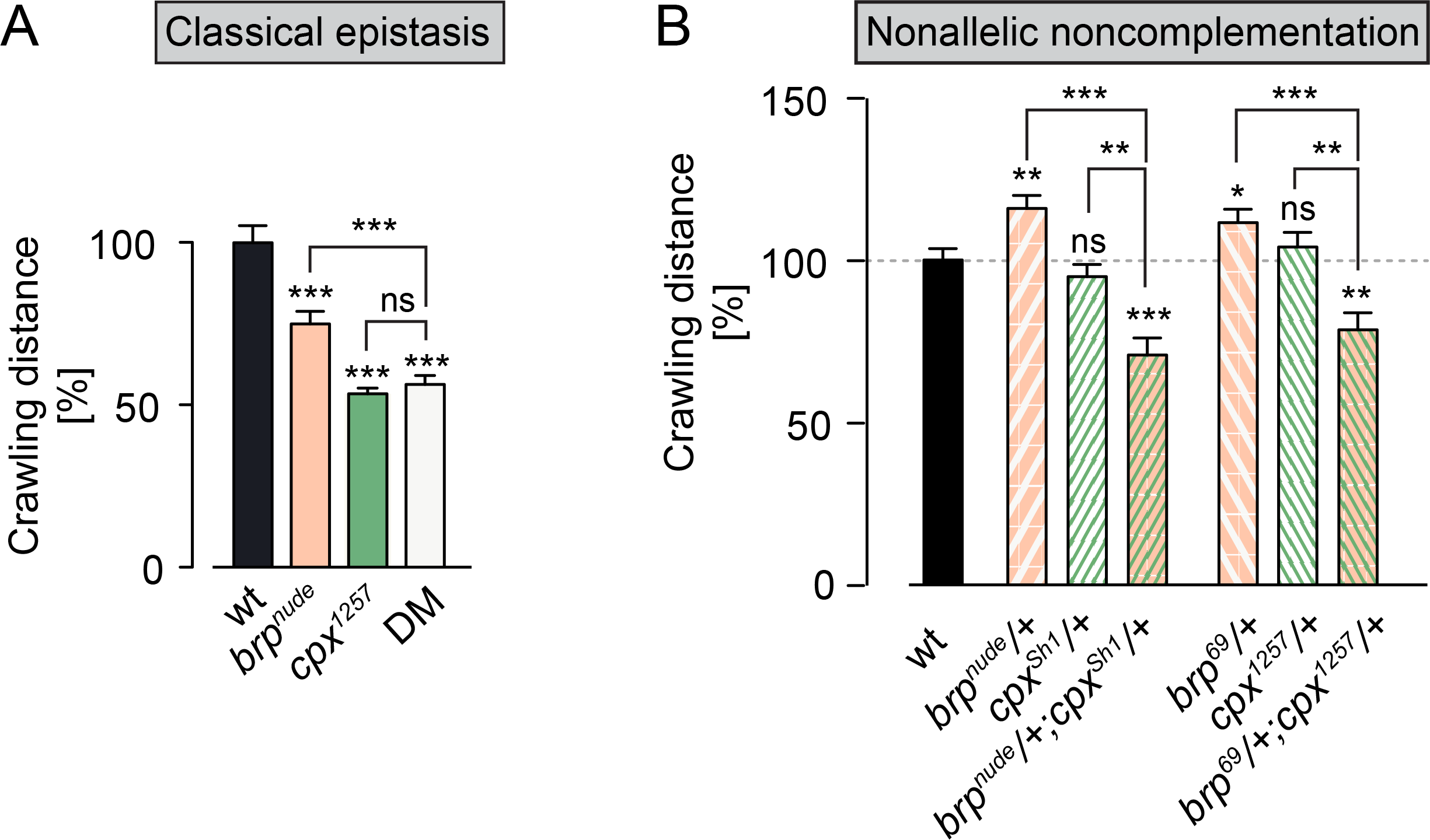
Genetic interaction between *cpx* and *brp*. **(A)** Quantification of crawling distances covered by larvae over a period of 2 min. The severe locomotion defects of *brp^nude^* (*brp^nude^/brp^69^*) and *cpx^1257^* (*cpx^1257^/cpx^Sh1^*) single mutants are not additive in the double mutant (DM). **(B)** Whereas larvae carrying one mutant copy of *cpx* or *brp* crawl normally, the double heterozygotes (*brp^nude^/+;cpx^Sh1^/+* and *brp^69^/+;cpx^1257^/+*) display impaired locomotion. This supports the notion that Brp and Cpx function in a common signalling pathway. Data presented as mean ± SEM, * P ≤ 0.05, ** P ≤ 0.01, *** P ≤ 0.001.

Nonallelic noncomplementation describes the failure of recessive mutations in two distinct loci to complement one another. Put differently, the double heterozygote displays a phenotype despite the presence of a wt copy of each gene. This phenomenon often indicates a physical interaction between the two gene products. Moreover, nonallelic noncomplementation assays in *C. elegans* have demonstrated that synaptic function is sensitive to hypomorphic mutations, whose altered gene products may ‘poison’ a limited number of presynaptic protein complexes (Yook et al., 2001). We therefore tested whether *brp^nude^* exhibited noncomplementation with a *cpx* null allele (*cpx^Sh1^*) and, vice versa, whether *cpx^1257^* exhibited noncomplementation with a *brp* null allele (*brp^69^).* Larval crawling measurements revealed impaired locomotion in the double heterozygotes, which were not observed in either heterozygote alone (**Figure 3B**, **Table S1**). These defects were synergistic and not additive effects of the noncomplementing mutations. Thus, *brp^nude^* and *cpx^1257^* mutations sensitize synaptic function to decreased levels of Cpx and Brp, respectively. These results further support the view that Brp and Cpx share a functional pathway.

### Disruption of Cpx targeting to SVs impairs their tethering to the CAZ

In electron microscopy (EM), the *Drosophila* CAZ is frequently associated with an electron-dense, T-shaped protrusion (termed T-bar) extending inwards from the active zone membrane, with Brp^C-tip^ marking its membrane-distal region (**Figure 4A**). As the *brp^nude^* mutant is characterized by a reduced number of SVs morphologically tethered to the T-bar (Hallermann et al., 2010b), we investigated whether the *cpx^1257^* mutation also led to decreased SV tethering by the T-bar. Quantifying the number of SVs within four concentric shells from 50-250 nm around the T-bar (**Figure 4B**, **Table S5**) revealed that both *brp^nude^* and *cpx^1257^* mutations impair SV tethering, leading to fewer SVs in the first three shells. Importantly, these morphological phenotypes were not additive in the double mutant. This strongly suggests that an interaction between Brp and Cpx helps tether SVs to the CAZ.

**Figure 4.**
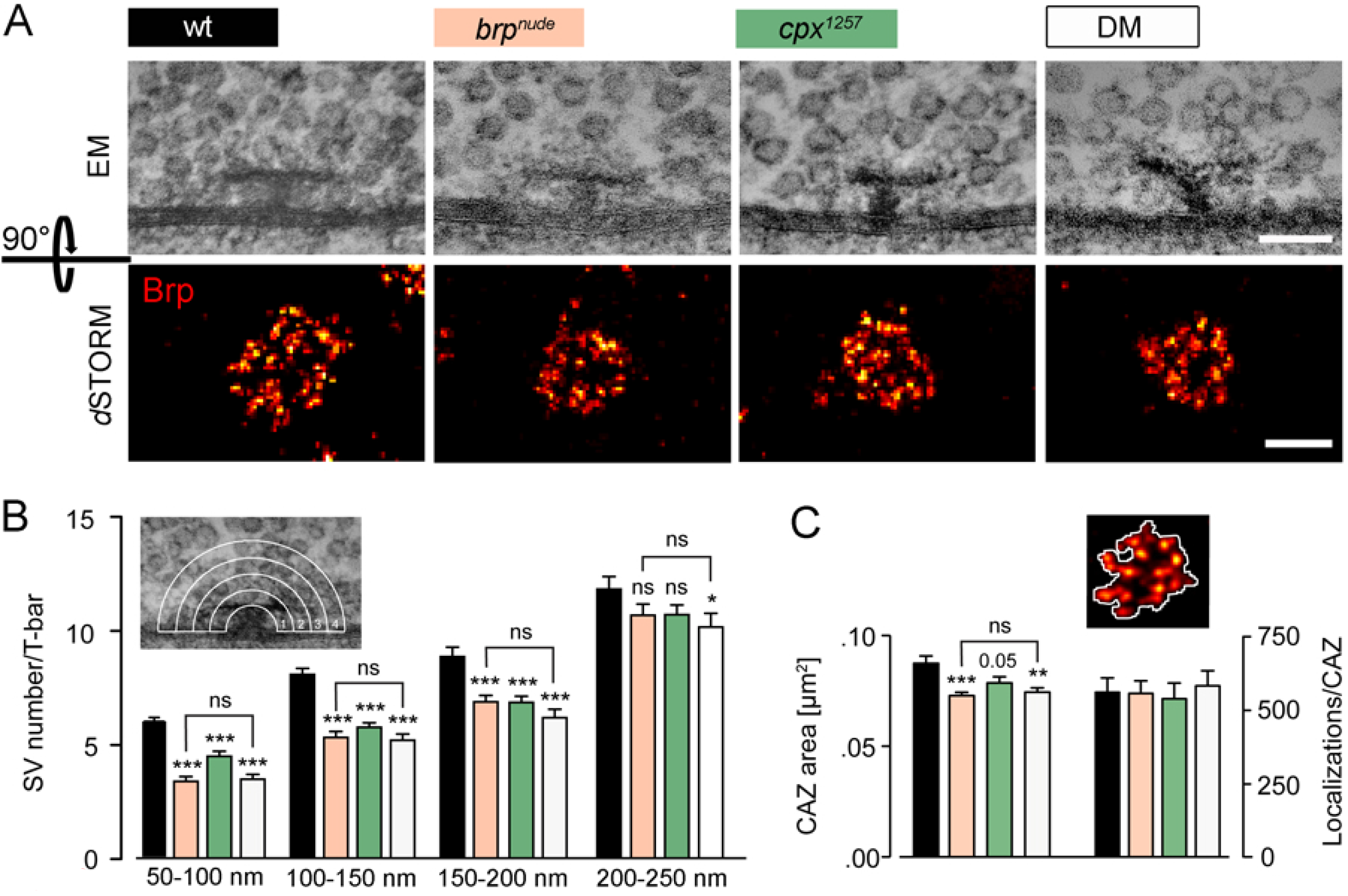
Cpx supports SV tethering to the *Drosophila* CAZ. **(A)** Upper panels, T-bars (viewed from the side) with associated SVs demark the CAZ in electron micrographs of larval motoneurons. Lower panels, *d*STORM images of antibody stainings against Brp [monoclonal antibody (mAb) nc82, NMJ 6/7] show the CAZ viewed *en face*. **(B)** Quantification of SVs surrounding the T-bar in four concentric shells, each 50 nm wide. Compared to the wt (black), fewer SVs are found near the T-bar in *brp^nude^* (beige) and *cpx^1257^* (green) single mutants. A comparable, non-additive effect is seen in the double mutant (DM, white). **(C)** The count of fluorophore localization events in *d*STORM images is proportional to the number of Brp protein copies (Ehmann et al., 2014). In all three mutant genotypes, the same number of Brp proteins occupies a smaller area. This is in line with CAZ filaments spreading out upon SV tethering. Data presented as mean ± SEM, * P ≤ 0.05, ** P ≤ 0.01, *** P ≤ 0.001 (vs. wt). Scale bars (A) upper panel 100 nm, lower panel 200 nm.

Using super-resolution fluorescence imaging by *direct* stochastic optical reconstruction microscopy (*d*STORM) (Heilemann et al., 2008; van de Linde et al., 2011), it was recently demonstrated that the diminished tethering capacity of the *brp^nude^* CAZ is accompanied by subtle modifications of its nanoarchitecture (Ehmann et al., 2014). An average CAZ-unit, equivalent to a T-bar, has been estimated to contain ~137 Brp proteins. Hence, antibody staining against this major CAZ component can provide detailed information on the CAZ ultrastructure when interrogated by *d*STORM with a spatial resolution of ~20 nm (Ehmann et al., 2014). Viewed *en face* (i.e. with the optical axis perpendicular to the active zone membrane), *d*STORM depicts the CAZ as a ring-like organization of Brp clusters, likely reflecting individual T-bar filaments. In *brp^nude^* mutants a normal number of Brp proteins are distributed over a smaller area (**Figure 4A, C**, **Table S6**) (Ehmann et al., 2014). This observation is consistent with a spreading out of T-bar filaments in the SV-tethered state. On the whole, the *cpx^1257^* mutant CAZ was normally structured but, interestingly, also appeared subtly compacted. This ultrastructural phenotype was not additive in the double mutant. Thus, these results are consistent with the EM data (compare **Figure 4B** and **C**) and the model that Brp and Cpx act in one pathway to tether SVs to the CAZ.

### Disruption of Cpx targeting to SVs enhances short-term synaptic depression

Next, we performed electrophysiological measurements to investigate synaptic function. As previously reported (Buhl et al. 2013; Iyer et al., 2013), C-terminal truncation of Cpx triggers a massive increase in the rate of spontaneous SV fusions, manifested as miniature excitatory postsynaptic currents (minis) in two-electrode voltage clamp (TEVC) recordings (**Figure 5A, B**, **Table S7**). At the *Drosophila* NMJ, this effect appears to be caused by impaired clamping of the fusion machinery when Cpx is not localized properly to exocytosis sites. In *cpx^1257^* mutants, the elevated mini frequency was accompanied by additional active zone formation (**Figure 5C, D**, **Table S8**). This matches the NMJ outgrowth observed under conditions of increased spontaneous neurotransmitter release (Choi et al., 2014). The mini frequency of *brp^nude^* mutants was comparable to wt. Surprisingly, however, the *brp^nude^* allele partially rescued the high mini frequency of *cpx^1257^* mutants and reverted the increased number of active zones (**Figure 5A-D**, **Tables S7**, **S8**). This highlights that Brp and Cpx perform distinct functions at the active zone. In contrast to Cpx, Brp^C-tip^ is not critical for clamping SVs in a release-ready state and loss of Brp^C-tip^ imposes a limit on the rate of spontaneous SV release through a Cpx-independent pathway.

**Figure 5.**
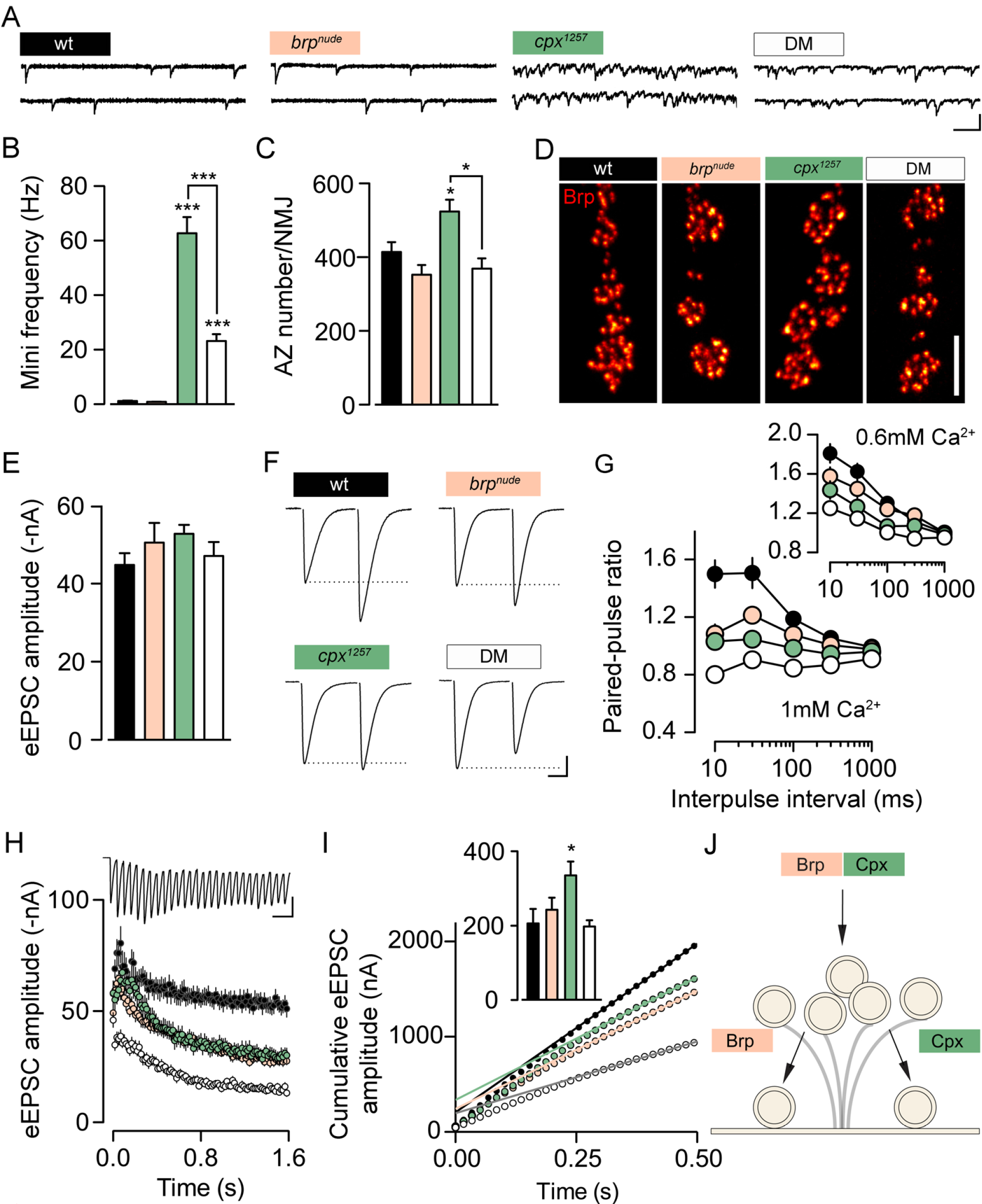
Brp and Cpx accelerate SV reloading through separate pathways. **(A)** Example traces of TEVC recordings from larval NMJs and **(B)** quantification of data show the dramatically increased mini frequency of the *cpx^1257^* mutant (green), which is partially rescued in the double mutant (DM, white). **(C)** Additional active zone formation accompanies the elevated mini frequency in the *cpx^1257^* mutant. **(D)** Shown are confocal images of Brp-stained NMJs (muscles 6/7). **(E)** The amplitude of eEPSCs is comparable in all genotypes at 0.2 Hz stimulation. **(F, G)** At short intervals, paired-pulse stimulation evokes less facilitation in the single mutants (*brp^nude^*, beige) than at wt synapses (black). This trend is further enhanced in the double mutant. **(H)** Average eEPSC amplitudes at 60 Hz stimulation and **(I)** cumulative amplitude plot. Back-extrapolation of a linear fit (0.3-0.5 s) yields an estimate of the RRV pool size (inset in nA). **(J)** Schematic working model: Brp and Cpx cooperate to tether SVs to the CAZ and function in parallel pathways to support SV recruitment to release sites. Unless noted otherwise, TEVC recordings were made in 1 mM [Ca^2+^]_e_. Data presented as mean ± SEM, * P ≤ 0.05, *** P ≤ 0.001. Scale bars (A) 100 ms, 3 nA, (D) 5 μm, (F) 10 ms, 10 nA, (H) 50 ms, 40 nA.

To obtain more information on synaptic physiology, we next analyzed eEPSCs. Despite normal basal current amplitudes at low frequency stimulation (0.2 Hz; **Figure 5E**, **Table S7**) both *brp^nude^* and *cpx^1257^* synapses showed disrupted short-term facilitation during paired-pulse stimulation (**Figure 5F, G**, **Table S7**) (Hallermann et al., 2010b). Remarkably, this trend was further increased at double mutant synapses, resulting in low facilitation in 0.6 mM extracellular Ca^2+^ concentration ([Ca^2+^]_e_) and depression in 1 mM [Ca^2+^]_e_ (**Figure 5F, G**, **Table S7**; rank sum test versus Control, 1mM [Ca^2+^]_e_: *brp^nude^*, 10 ms P = 0.0039, 30 ms P = 0.0101, 100ms P = 0.02, all other intervals n.s.; *cpx^1257^*, 1000 ms n.s., all other intervals P ≤ 0.001; DM, 10 ms P = 0.002, 1000 ms P = 0.0011, all other intervals P ≤ 0.0001; 0.6mM [Ca^2+^]_e_: *brp^nude^*, all intervals n.s.; *cpx^1257^*, 10 ms P = 0.0123, 30 ms P = 0.0019, 100 ms P = 0.0026, all other intervals n.s.; DM, 1000 ms n.s., all other intervals P ≤ 0.001). These observations point towards an additive effect of *brp^nude^* and *cpx^1257^* mutations on short-term depression, which we investigated in more detail.

Desensitization of postsynaptic glutamate receptors shapes the decay time constant (τ_decay_) of eEPSCs and can contribute to use-dependent depression at the *Drosophila* NMJ (DiAntonio et al., 1999; Heckmann and Dudel, 1997; Schmid et al., 2008). Since the τ_decay_ was not shortened in any of the mutant genotypes, enhanced receptor desensitization was likely not the cause of impaired facilitation (**Figure S5**, **Table S7**). Presynaptic alterations that promote short-term depression without changing basal eEPSC amplitudes include slowed SV recruitment, as in the case of *brp^nude^* mutants, or an elevated release probability of fewer RRVs (Ehmann et al., 2014). In both cases, normal SV release may be maintained during low frequency stimulation, but as stimulation intervals shorten, SV availability becomes rate limiting and compound release drops. In order to differentiate between these two sources of depression, we estimated the size of the RRV pool in the different genotypes. Consistent with the paired-pulse protocols, a train of high frequency stimulation (60 Hz) provoked similar short-term depression of eEPSC amplitudes at *brp^nude^* and *cpx^1257^* synapses and triggered enhanced depression in the double mutant (**Figure 5H**). Pool estimates can be derived by back-extrapolation of a linear fit to the late phase of cumulatively plotted eEPSCs (Schneggenburger et al., 1999). While the quantitative interpretation of extrapolation-based pool estimates must be treated with caution (Hallermann et al., 2010a; Neher, 2015), we found no evidence of fewer RRVs in the single mutants, nor in the double mutant (**Figures 5I**, **S6**, **Table S9**). Thus, these results are consistent with slow SV reloading at *brp^nude^* and *cpx^1257^* active zones. Moreover, the additivity of electrophysiological phenotypes in the double mutant suggests that Brp- and Cpx-dependent SV replenishment operates through separate mechanisms.

The altered short-term plasticity of *cpx^1257^* mutants prompted us to interrogate the molecular organization of their active zones in more detail. Imaging by Structured Illumination Microscopy (SIM; ~2-fold increase in 3-D resolution compared to conventional wide-field microscopy) (Gustafsson et al., 2008) revealed an increased number of Ca^2+^-channel clusters at *cpx^1257^* NMJs (**Figure S7**, **Table S10**) proportional to the elevated number of active zones (**Figure 5C**). Interestingly, whereas the arrangement of active zone Ca^2+^-channels is unaffected by the *brp^nude^* mutation (Hallermann et al., 2010b), the average cluster size is slightly reduced in *cpx^1257^* mutants (**Figure S7**, **Table S10**). This might give rise to a diminished release probability, which in turn could explain why *cpx^1257^* NMJs display normal basal EPSC amplitudes despite possessing more release sites (**Figure 5C**, **Table S8**).

Taken together, our results support a model whereby Brp and Cpx function in the same pathway to tether SVs to the CAZ (no additivity of ultrastructural phenotypes; **Figure 4**), but act in parallel pathways to promote rapid recruitment of tethered SVs to the active zone membrane (additivity of short-term depression; **Figure 5J**).

### SV tethering by Cpx is evolutionarily conserved

To test whether SV tethering is a specialization of *Drosophila* Cpx or an evolutionarily conserved property, we turned to rod bipolar cells (RBs) of the mouse retina. RB synapses onto AII amacrine cells are glutamatergic, they possess prominent electron-dense ribbons as CAZ specializations and express only Cpx3, a retina-specific isoform. Like the main isoform at the *Drosophila* NMJ, mammalian Cpx3 is also farnesylated at a C-terminal CAAX motif (Reim et al., 2005). We re-analyzed ultrastructural data from (Mortensen et al., 2016) and focused specifically on the number of ribbon-proximal SVs in electron micrographs of RB terminals. In *cpx3* knockout mutants (*cpx3^KO^*) significantly fewer SVs were found in concentric shells from 0-400 nm around the ribbon (**Figure 6**, **Table S11**), despite a normal cytoplasmic area covered by the shells (**Figure S8**, **Table S11**). This result resembles the morphological defect of *cpx^1257^* mutants at the *Drosophila* NMJ and supports the notion that SV tethering is an evolutionarily conserved feature of certain Cpx family members.

**Figure 6.**
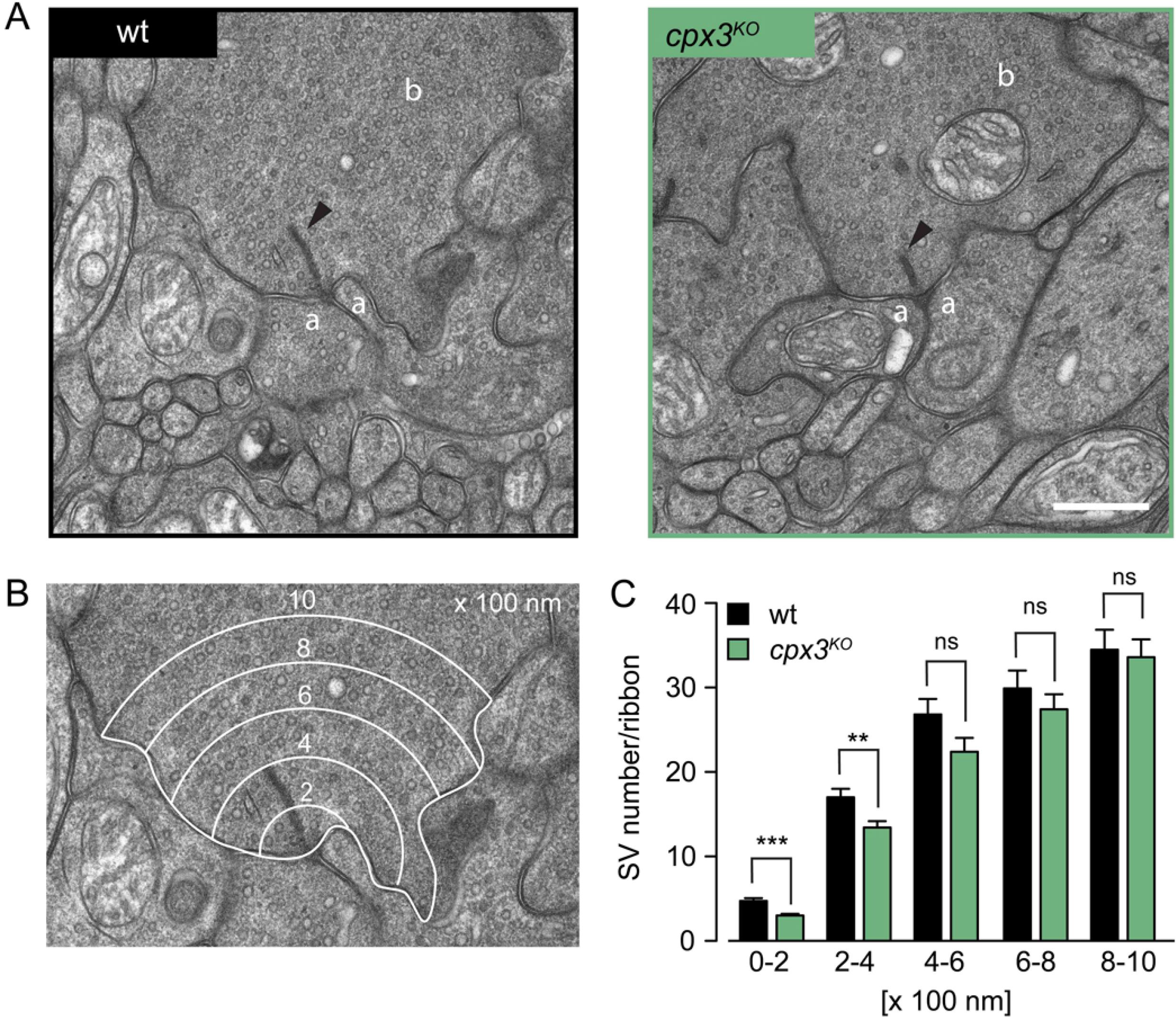
Mouse Cpx3 promotes SV tethering to ribbons of rod bipolar cells. **(A)** Electron micrographs show wt (left) and *cpx3^KO^* (right) ribbon synapses of mouse rod bipolar cells. The arrowheads indicate electron dense ribbons to which SVs are tethered. Amacrine cell process (a), RB terminal (b). **(B)** Example of an electron micrograph overlaid with five concentric shells, each 200 nm wide (0-200 nm; 200-400 nm, 400-600 nm, 600-800 nm and 800-1000 nm), centered at the ribbon base. **(C)** Counting the SVs in each shell revealed a significant drop of SVs within 0-400 nm around *cpx3^KO^* ribbons. Data presented as mean ± SEM, ** P ≤ 0.01, *** P ≤ 0.001. Scale bar 500 nm.

## DISCUSSION

Tethering SVs to fine filamentous CAZ structures is believed to support their rapid reloading at the active zone membrane to sustain high neurotransmitter release rates (Hallermann and Silver, 2013). Correspondingly, disrupting CAZ tethers impedes ongoing SV release (Frank et al., 2010; Hallermann et al., 2010b, 2010c; Miskiewicz et al., 2014; Snellman et al., 2011). The experiments reported here identify Cpx as a Brp interactor, which promotes SV tethering to the CAZ and counteracts short-term synaptic depression. Similarly, recent work on SV recycling at the *Drosophila* NMJ also found a reduced number of T-bar associated SVs in *cpx^null^* mutants (Sabeva et al., 2017). Cpx has been studied extensively in its capacity as a SNARE accessory protein, which regulates the fusion complex (Brose, 2008; Neher, 2010a; Reim, 2017; Trimbuch and Rosenmund, 2016). By interacting with the assembled trans-SNARE complex, Cpx promotes Ca^2+^-triggered evoked neurotransmitter release and, depending on species, synapse type and preparation, either facilitates or clamps spontaneous release. At *C. elegans* and *Drosophila* NMJs, loss of Cpx leads to a dramatic increase in mini frequency (Huntwork and Littleton, 2007; Martin et al., 2011; Xue et al., 2009) (**Figure 5A, B**, **Table S7**), highlighting the bipolar function of Cpx at these invertebrate active zones. Here, Cpx appears to maintain SVs in a release-competent state by both blocking fusion and by supporting priming (Hobson et al., 2011). Our results indicate that the release-promoting role of Cpx extends even further upstream to facilitate SV recruitment to CAZ filaments and to the active zone membrane. This suggests that Cpx links the final stages of the SV cycle preceding exocytosis.

The hypomorphic *cpx^1257^* allele strongly decreases Cpx protein levels and changes its localization pattern at the NMJ (**Figures 2D**, **S3**, **S4**), consistent with previous reports linking C-terminal farnesylation of Cpx to synapse targeting (Buhl et al. 2013; Iyer et al., 2013). Despite maintaining normal basal eEPSC amplitudes, the hypomorphic mutants display a roughly 60-fold increase in mini frequency (**Figure 5A, B**, **Table S7**). The different susceptibility of evoked and spontaneous fusion modes to reduced Cpx levels supports the notion that the clamping role of Cpx dominates at the *Drosophila* NMJ (Huntwork and Littleton, 2007; Iyer et al., 2013; Xue et al., 2009). Why does the C-terminal Brp truncation partially rescue the clamping defect of *cpx^1257^* active zones? One explanation could be that this phenomenon is directly related to the additivity of short-term depression in the double mutant (**Figure 5H**, **Table S7**). If Cpx and Brp function in parallel to promote reloading of tethered SVs to the active zone membrane, then removing both pathways could limit both the maximal rate of evoked and (unclamped) spontaneous transmitter release. In *brp^nude^* single mutants, in turn, the intact Cpx clamp would prevent slowed reloading from reducing the mini frequency. Alternatively, evoked and spontaneous transmission may use nonidentical SV pools regulated differentially at the molecular level (Crawford and Kavalali, 2015; Kaeser and Regehr, 2014; Melom et al., 2013).

What could the parallel Cpx- and Brp-dependent pathways of SV replenishment correspond to mechanistically? It is tempting to speculate that Cpx may facilitate SV recruitment to activated release sites by helping to bring v-SNAREs and t-SNAREs within striking distance of each other to initiate fusion complex formation (Zenisek et al., 2000). On the other hand, Cpx may assist in release site activation itself, i.e. removal of SV proteins from prior fusion events and resetting of the release machinery. Given a limited number of release sites, their delayed restoration will enhance short-term depression (Hosoi et al., 2009; Kawasaki et al., 2000; Neher, 2010b). Similarly, Brp filaments may guide SVs to release sites or assist in site-clearance. Therefore, an attractive model that describes parallel functional pathways for Brp and Cpx (producing additive short-term depression phenotypes in the double mutant) is that one of the two accelerates SV reloading and the other promotes release site activation.

Our morphological data point towards an interaction between Cpx and Brp in tethering SVs to the CAZ. Thus, defective tethering should contribute similarly to the functional phenotypes seen in both single mutants. SV tethering may enhance docking efficiency and rapid exocytosis by restricting the diffusion of SVs and thereby concentrating them in the vicinity of release sites. Consistent with this notion, findings at several synapses have suggested that undocked SVs located near the active zone membrane can be recruited and released very rapidly (Hallermann et al., 2010b; Jockusch et al., 2007; Rizzoli and Betz, 2004; Wang et al., 2016). Our results suggest that Cpx associates with SVs (**Figures 2D**, **S3**, **S4**) and indicate that this interaction serves a dual function: it concentrates Cpx at synaptic sites and facilitates SV binding to Brp filaments. The association of Cpx with SVs has previously been reported in motoneurons of *C. elegans*, as well as larval and adult *Drosophila*. Whereas SV targeting and synaptic localization of Cpx depend on a C-terminal amphiphatic region in *C. elegans*, the predominant *Drosophila* isoform appears to associate with SVs through a membrane-binding farnesylated CAAX motif at its C-terminus (Buhl et al. 2013; Cho et al., 2010; Iyer et al., 2013; Wragg et al., 2013; Xue et al., 2009). Mammalian Cpx isoforms 3 and 4 are also C-terminally farnesylated (unlike Cpx1 and 2) and here, too, this posttranslational modification is required for Cpx targeting to synapses (Reim et al., 2005, 2009). *Drosophila* Cpx is more mobile than SVs and less mobile than cytoplasmic GFP, indicating a low-affinity transient interaction with Cpx cycling on and off SVs (Buhl et al. 2013). Similarly, biochemically purified SVs from rat brain show that mammalian Cpx1 and 2 do not bind SVs with high affinity (Denker et al., 2011; Wilhelm et al., 2014) though the curvature-sensing C-terminal domain appears to preferentially localize Cpx1 to SV membranes (Gong et al., 2016). A transient association between Cpx and Brp would explain why Cpx was not pulled down by immobilized Brp^C-tip^ in a biochemical analysis (**Figure S9**). In this case, our in vivo setting was likely necessary to identify this functional protein interaction. Alternatively, an extended C-terminal fragment of Brp (Brp^C-long^) may be necessary to capture Cpx in vitro, similar to its requirement for ectopic axonal SV enrichment (**Figure 1G**).

It is conceivable that our in vivo screen may have scored false negatives. That is, further SV proteins likely participate in CAZ tethering and some of these may have passed undetected due to e.g. inefficient RNAi knock-down. Moreover, it should be noted that additional Brp domains and/or other CAZ proteins must be involved in tethering, since a physical association of SVs with the CAZ still persists in *brp^nude^* mutants, albeit at lower levels (**Figure 4**, **Table S5**) (Hallermann et al., 2010b).

Of the four mammalian *cpx* genes, *cpx1* and *2* are the main isoforms in the central nervous system. *cpx3* and *4*, in turn, are predominantly expressed in the retina where they are specifically localized to ribbon synapses of photoreceptors and bipolar cells (Reim et al., 2005, 2009). Strikingly, knockout of *cpx3* significantly reduced SV tethering to the RB ribbon (**Figure 6**, **Table S11**), highlighting intriguing parallels between this synapse and the *Drosophila* NMJ. In both glutamatergic neurons, C-terminal farnesylation of Cpx is required for its synaptic localization and at both active zones, Cpx participates in SV tethering to prominent electron-dense specializations of the CAZ (T-bar and ribbon at the NMJ and RB, respectively). This could indicate a specialized tethering function of farnesylated Cpx at ribbon-like CAZ structures. Alternatively, SV tethering by Cpx may be widespread in the vertebrate CNS and is simply more easily detected at ribbon synapses, where the association of SVs with the CAZ is very prominent. Either way, our findings argue that Cpx-dependent SV tethering is an evolutionarily conserved property of this multifunctional regulator of active zone function.

## ACKNOWLEDGEMENTS

We thank T. Littleton for sharing the Cpx antibody, F. Kawasaki for *cpx^1257^* flies, L. van Werven, Thea Hellmann, M. Oppmann and B. Trost for expert technical assistance, C. Wichmann for ultrastructural advice, as well as E. Buchner and S. Hallermann for discussions and M. Sauer for supporting SIM experiments. This work was funded by grants from the Deutsche Forschungsgemeinschaft to T.L. (FOR 2149/P01 and P03, SFB 1047/A05, TRR 166/C03, LA2861/7-1), R.J.K. (FOR 2149/P03, SFB 1047/A05, TRR 166/B04, KI1460/4-1) and N.S. (FOR 2149/P01). N.E. was supported by a grant from the German Excellence Initiative to the GSLS, University of Würzburg, GSC106/3 (PostDoc Plus program) and N.S. was supported by a Junior Research grant from the Medical Faculty, University Leipzig. The authors declare no conflict of interest.

## MATERIALS AND METHODS

### *Drosophila* culture and stocks

*Drosophila* were cultured on standard cornmeal food at 25 °C. The following strains were generated in this study:

TAG61, *w^1118^; P{UAS-3xFlag::amCFP::brp^C17^ w^+^}/CyO;;* (BRP^C-tip^)
TAG126, *w^1118^; P{UAS-3xFlag::amCFP w^+^}/Cyo;;* (CFP)
TAG77, *w^1118^;; P{UAS-mCD8::EGFP w^+^}^attP2^/TM3, Sb;* (CD8::EGFP = no BRP)
TAG70, *w^1118^;; P{UAS-mCD8::EGFP::brp^C17^ w^+^}^attP2^/TM3, Sb;* (CD8::EGFP::BRP^C-tip^)
TAG118, *w^1118^;; P{UAS-mCD8::EGFP::brp^C862^ w^+^}^attP2^/TM3, Sb;* (CD8::EGFP::BRP^C-long^)
TAG151, *w^1118^; P{UAS-mRFP::syt-1 w^+^}^attP40^/Cyo;;* (mRFP::SYT-1)
*w^*^; P{UAS-cac1::EGFP} ok6-GAL4 w^+^/CyOGFP W; cpx^sh1^/TM6b, Tb;*

Additional strains are listed below. Flies were obtained from the Bloomington Stock Center (NIH P40OD018537), the Harvard Exelixis deficiency collection, the Vienna *Drosophila* Research Center (RNAi strains) (Dietzl et al., 2007), or were gifts from colleagues.

*w^1118^; brp^nude^/CyoactGFP w^+^;;* (Hallermann et al., 2010b)
*w^1118^; brp^69^/CyoGFP w^−^;;* (Kittel et al., 2006)
*w^*^;; cpx^sh1^/TM6b, Tb;* (Huntwork and Littleton, 2007)
*w^*^;; cpx^1257^/TM6c* (Iyer et al., 2013)
*w^*^; ok6-GAL4 w^+^;;* (Sanyal, 2009)
*w^*^; Vglut-GAL4 w^+^;;* (Daniels et al., 2008)
*w^*^;; Actin-5C-GAL4 w^+^;* (Ito et al., 1997)
*w^*^; P{UAS-cac1::EGFP} ok6-GAL4 w^+^/CyOGFP w;;* (Kawasaki et al., 2004)
*w^1118^;; Df(3R)Exel6140/TM6b, Tb;* (gift from F. Kawasaki, BL# 7619)
*w^*^;; P{GD10482}v21477; (cpx^RNAi^)*
*w^*^; P{GD9877}v33606;; (dunc13^RNAi^)*
*w^*^;; P{GD8641}v43629; (tomosyn^RNAi^)*
*w^*^; P{GD14785}v29969;; (dCirl^RNAi^)*
*w^*^; P{GD9502}v25291;; (dCAPS^RNAi^)*
*w^*^; P{KK109431}v103201;; (csp^RNAi^)*
*w^*^; P{GD 1171}v47506;; (syt-12^RNAi^)*
*w^*^;; P{GD8644}v24988; (syt-7^RNAi^)*
*w^*^; P{GD15273}v39384;; (rim-1^RNAi^)*
*w^*^;; P{GD12118}v27824/TM3; (rab3-GAP^RNAi^)*
*w^*^;; P{GD11312}v26537/TM3; (gdi^RNAi^)*
*w^*^; P{GD7330}v52438;; (rph^RNAi^)*
*w^1118^ P{GD17411}v49247;;; (rac-1^RNAi^)*
*w^*^;; P{GD10492}v34094; (rab5^RNAi^)*
*w^*^; P{GD3785}v8784;; (sng-1^RNAi^)*
*w^*^;; P{KK108941}VIE-260B; v110606 (syn^RNAi^)*
*w^*^;; P{GD11750}v27493 (annb9^RNAi^)*
*w^*^;; P{GD14255}v36107; (annb10^RNAi^)*
*w^*^;; P{GD10342}v25817; (twf^RNAi^)*
*w^*^;; P{GD10754}v34354; (dysb^RNAi^)*
*w^*^;; P{GD7149}v16133; (atg-1^RNAi^)*
*w^*^; P{GD 1254}v5702;; (rsk^RNAi^)*
*w^*^; P{GD12534}v35445;; (sap47^RNAi^)*
*w^*^;; P{GD12383}v35346; (dsyd-1^RNAi^)*
*w^*^; P{GD834}v2574;; (dvglut^RNAi^)*
*w^*^; P{GD2233}v45726;; (vti1a^RNAi^)*
*w^1118^; P{GD2076}v10061/TM3; (mctp^RNAi^)*

### Transgene construction

#### pNH28 (BRP^C-tip^)

amCFP was amplified from *pTL304* using *nh_56F/57R* and cloned into *pENTR™3C* (Invitrogen, Cat.No.: A10464) with *KpnI* and *XhoI* (pNH27). *nh_57R* contained the last C-terminal 17 amino acids of Brp. LR-based recombination of *pNH27* with expression plasmid TFW provided the N-terminal *3xFlag* and led to *pNH28*.

#### pNH54 (CFP)

Primers *nh_56F/75R* were used to amplify amCFP from *pNH28*. The resulting *XhoI/KpnI*-containing fragment was cloned into *pENTR™3C* (*pNH53*). Subsequently, an LR reaction with *pTFW* resulted in *pNH54*.

#### pNH42 (no BRP)

The *mCD8::EGFP* fusion construct was amplified from *pTL231* using primers *tl_330F* and *nh_02R*. The resulting 1.4 kb fragment, which carried a Kozak sequence was cloned into *pENTR™3C* using *DraI* and *XhoI* (*pNH32*). Next, the insert was recombined into *pTW-attB* using an LR recombination reaction, which yielded the expression construct *pNH42*.

#### PNH39 (BRP^C-tip^)

The *mCD8::EGFP* fusion construct was amplified from *pTL231*. Primers *tl_330F* and *nh_3R* were used to add a Kozak sequence, a 5x glycine linker and a sequence encoding the last C-terminal 17 aa of Brp. Amplicon insertion into *pENTR™3C* was done using *DraI* and *XhoI* and resulted in *pNH33*. LR recombination into *pTW-attB* led to the expression construct *pNH39*.

#### pNH52 (BRP^C-long^)

A 2.6 kb fragment encoding the last C-terminal 862 amino acids of Brp was amplified from *pTL319* with primers *nh_25F/24R*. The amplicon was digested with *DraI* and *XbaI* and subsequently ligated with *pNH43*, which resulted in the ENTRY clone *pNH51* and after recombination in the expression construct *pNH52*.

#### pNH60 (mRFP::SYT)

The amplification of *syt* from *pTL143* was performed using primers *nh_94F/95R*. mRFP was amplified from *pNH11* using *nh_96F/97R*. Both PCR fragments were cloned into *pTW-attB* using *XhoI, NheI* and *AgeI* via triple ligation.

The AccuStar high-fidelity proof-reading DNA polymerase (Eurogentec) was utilized for all PCR-based cloning steps. Initial insert verification was done by restriction analyses. To ensure absence of errors each PCR-amplified region was completely sequenced.

#### Primers (5’-3’ orientation)

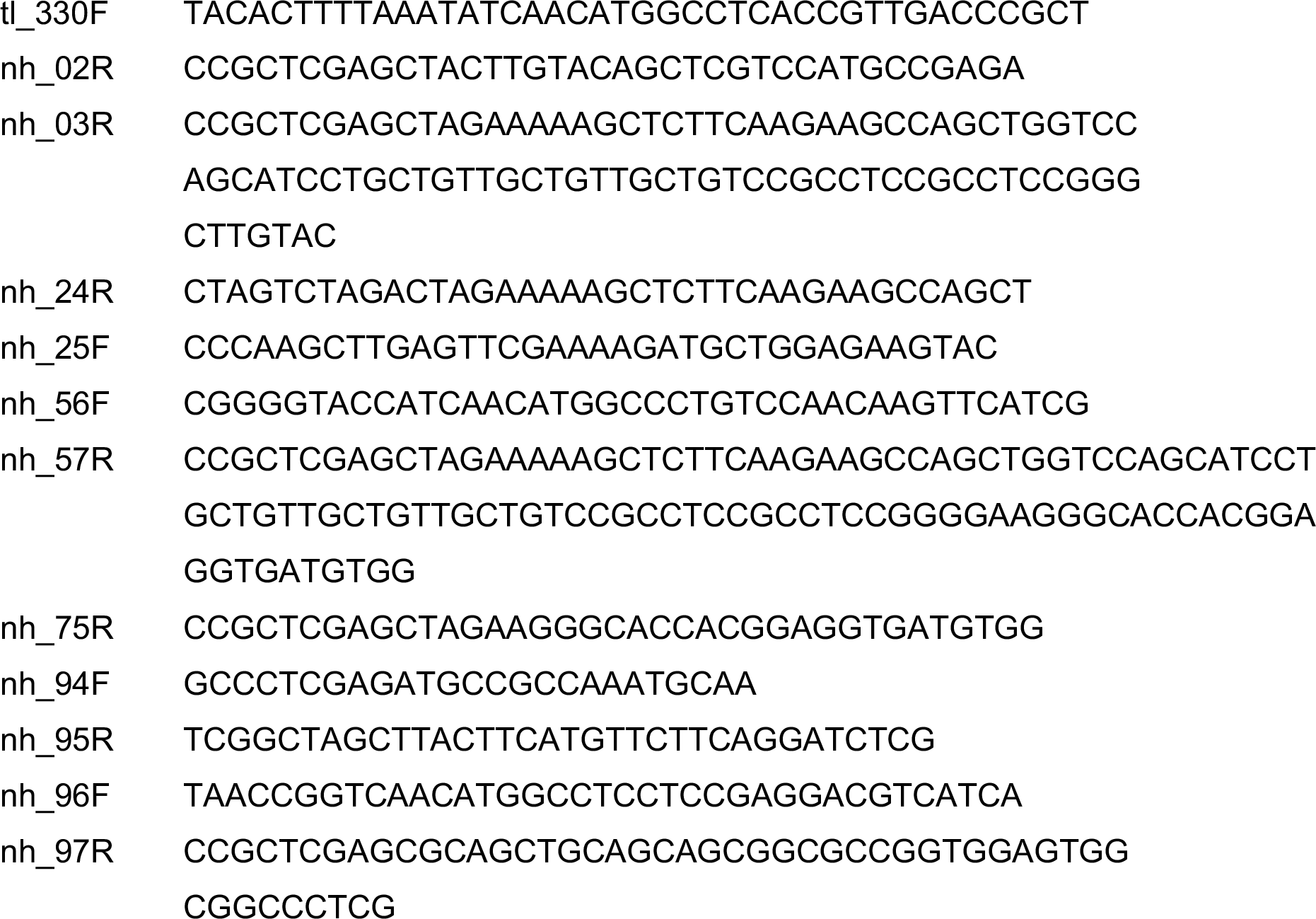

### PhiC31-mediated integration of transgenes

Germ-line transformation of transgenes was done by BestGene Inc. either by random P-element transformation or site-specific phiC31 methodology (Groth et al., 2004). The vectors utilized were w^+^-marked allowing w^+^-based selection of recombinants. Specific integration sites attP40 (cyto site 25C6) (Markstein et al., 2008) and attP2 (BDSC stock# 8622) (Groth et al., 2004) were used for transgene insertion on second and third chromosomes, respectively. Brp bait variants (*CD8::EGFP, brp^C-tip^* and *brp^C-long^*) were integrated into attP2 to ensure similar expression conditions.

### Confocal microscopy

Wandering third instar *Drosophila* larvae were dissected in ice-cold HL-3 (hemolymph like solution) (Stewart et al., 1994), fixed for 10 min using 4% paraformaldehyde in 0.1 M phosphate buffered saline (PBS) and blocked for 30 min in PBT (PBS with 0.05% Triton X-100, Sigma-Aldrich), containing 5% normal goat serum (NGS, Jackson ImmunoResearch). Samples were incubated with primary antibodies at 4°C overnight. The next day, filets were incubated with secondary antibodies for 2 h at room temperature (RT). Each antibody incubation step was followed by 2 short and 3 × 20 min washing steps. Preparations were stored in Vectashield (Vector Laboratories) overnight at 4°C before mounting. Confocal recordings were performed with a Zeiss LSM5 Pascal confocal system. To ascertain comparability of different genotypes, larvae were stained in the same vial and NMJs on muscles 6/7 (segments A2 and A3) were imaged in one session. Also, image acquisition of genotypes alternated and was performed with identical laser settings if not stated otherwise. Image analysis was done using ImageJ (National Institutes of Health, Bethesda). Prior to image analysis, unspecific signals were manually removed from maximal projections of confocal stacks. To analyse AZ numbers, the background was subtracted from the maximal projections. A threshold (mean grey value 25) was set and a Gaussian blur (σ = 1 pixel) was applied to generate masks, which were subsequently superimposed onto the maximal projection. AZs were detected with the find ‘Maxima’ command and quantified via ‘Analyze Particles’.

To quantify SV levels, the mRFP::SYT signal was amplified using a polyclonal antiserum that recognizes RFP. For each larva, images were acquired at NMJ 6/7 (segment A2 or A3) and the nerve exiting the VNC at segment A8 (caudal end of the VNC). Laser settings were adjusted according to SV levels at the NMJ and subsequently retained to capture SV abundance in the nerve. This procedure was used to calculate the mRFP::SYT ratio of an individual larva and controlled for differences in expression strength across samples. After subtracting the background from maximal projections of confocal stacks, non-synaptic signals were manually removed from NMJ images, a minimal threshold was set (individually adjusted for each sample), a Gaussian blur (σ = 1 pixel) was applied and the average intensity of mRFP::SYT was measured. In the nerve, the mean mRFP::SYT signal intensity was measured several times within an area of defined size (54.1 μm^2^). The brightest region was used to calculate the NMJ/nerve ratio of SVs.

Antibodies were used in the following dilutions: mouse-α-Brp (nc82, 1:250; provided by E. Buchner; RRID: AB_528108) (Wagh et al., 2006), mouse-α-Csp (1:100; ab49, Developmental Studies Hybridoma Bank, RRID: AB_2307345), rabbit-α-Cpx (1:500; RRID: AB_2568068) (Huntwork and Littleton, 2007), rabbit-α-Vglut^N-term^ (1:500; RRID: AB_2567386, provided by A. DiAntonio) (Daniels et al., 2008), mouse-α-GFP (1:500, Sigma-Aldrich, RRID: AB_259941), rabbit-α-RFP (1:500, Antibodies-Online, RRID:AB_10781500), mouse-α-Flag (1:500; Sigma-Aldrich, RRID: AB_262044), α-Hrp conjugated with Cy3 (1:250; Jackson ImmunoResearch, RRID: AB 2338959), Alexa Fluor-488-conjugated goat-α-mouse (RRID: AB2534069) and goat-α-rabbit (RRID: AB_143165, both 1:250; Invitrogen Life Technologies), Cy3-conjugated goat-α-rabbit (RRID: AB_2338000) and goat-α-mouse antibodies (RRID: AB_2338690, both 1:250, Jackson ImmunoResearch).

### Super Resolution Microscopy

#### SIM

In principle, sample preparation followed the procedures described for confocal imaging. Control and mutant larvae were stained in the same vial and imaged in one session with the same laser settings to enable comparability. Images were acquired at NMJ 4, segments A2 and A3, with a Zeiss Elyra S.1 equipped with an oil immersion objective (Plan-Apochromat 63x/1.4) and 488 nm and 641 nm lasers. The Z step size was set to 0.2 μm and imaging was performed using 5 rotations of the grating at 5 different phase stepping. The raw images were processed with Zen imaging software (Carl Zeiss) and converted to lsm files. Maximal projections of optical slices were used for quantification. The analysis was carried out in ImageJ (National Institutes of Health) as described for confocal images using appropriate threshold and sizing criteria for Brp and Cacophony signals and genotypes were blinded to the investigator.

The following antibodies were used: mouse-α-Brp (nc82; 1:100), GFP-Booster ATTO647N (1:100; ChromoTek, RRID: AB_2629215) and Alexa Fluor-488-conjugated goat-α-mouse (1:250).

#### dSTORM

Super resolution imaging via *d*STORM was performed as previously described (Ehmann et al., 2014). Mouse-α-Brp (nc82) was used at a concentration of 1:2000 to quantify CAZ size and Brp localizations within the CAZ. Secondary goat-α-mouse F(ab’)_2_ fragments (A10534, Invitrogen) were labelled with Cy5-NHS (PA15101, GE Healthcare) yielding a degree of labelling of 1-1.3 and were used at a concentration of 5.2 × 10^−8^ M. Prior to *d*STORM recordings larval filets were embedded in photoswitching buffer, i.e. 100 mM mercaptoethylamine, pH 8.0, enzymatic oxygen scavenger system [5% (wt/vol) glucose, 5 U ml^−1^ glucose oxidase and 100 U ml^−^1 catalase] (Schäfer et al., 2013) and arranged on an inverted microscope (Olympus IX-71) equipped with an oil (60x, NA 1.49) or water (63x, 1.15) immersion objective and a nosepiece stage (Olympus IX2-NPS) (van de Linde et al., 2011). Positioning of filters and mirrors on a translational stage enabled switching between wide-field and low-angle/highly inclined thin illumination (van de Linde et al., 2011; Sharonov and Hochstrasser, 2007; Tokunaga et al., 2008). Optical components for excitation and emission of Cy5 are described elsewhere (Ehmann et al., 2014; Paul et al., 2015). Final pixel size was 126 nm (oil immersion objective) or 109 nm (water immersion objective). Data were reconstructed using rapi*d*STORM (Wolter et al., 2010, 2012). A sub-pixel binning of 10 nm was applied and fluorescence spots yielding >1000 photons were included in the analysis. To quantify CAZ size and Brp localizations within the CAZ, masks were created (Gaussian blur σ = 1 pixel), minimally thresholded (0.15 counts) and overlaid with the original image. Individual CAZs from NMJ 6/7 (segments A2 and A3) were identified via their area (300 pixel-infinity). Image analysis was performed using ImageJ. All genotypes were stained in the same vial and imaged in two sessions. To assure comparative imaging settings, data were analyzed for unspecific background labelling and excluded from the analysis if the background exceeded 2.3 single spots per μm^2^.

### Electron microscopy

#### Drosophila NMJ

Wandering third instar larvae were dissected in ice-cold HL-3 (Stewart et al., 1994) and fixed with glutaraldehyde solution [2.5% glutaraldehyde, 50 mM cacodylate buffer (pH 7.2), 50 mM KCl, 2.5 mM MgCl_2_, ddH_2_O] for 45 min at 25°C. Subsequently, samples were rinsed and washed 5 × for 3 min with 50 mM cacodylate buffer. Next, samples were fixed for 90-120 min with 2% OsO_4_ in 50 mM cacodylate buffer, shifted to aqueous media by washing 5 × in short intervals in distilled water. Then samples were contrasted overnight in aqueous 0.5% uranyl acetate. After 5 washes with ddH_2_O the samples were dehydrated in an ethanol series of 50%, 70%, 90%, 96% and 3 × 100% for 15 min each. In preparation of tissue embedding, samples were washed 2 × for 20 min with propylene oxide. Thereafter, the tissue was carefully infiltrated and embedded in conventional Epon, closed with a gelatine cap and cured at 60°C for 48 h. Subsequently, 60-80 nm ultra-thin sections were contrasted for 8 min using 5% filtered uranyl acetate in ethanol. In between and after incubation steps, sections were dipped into 100% ethanol, 50% ethanol and ddH_2_O. After careful drying, the sections were again contrasted with Reynolds’ lead citrate (for 4 min) (Reynolds, 1963) in decocted ddH_2_O and washed 3x in decocted ddH_2_O. Image acquisition was performed with a Zeiss EM 900. Images were registered on photo plates. The negatives were scanned (1200 dpi) to digitalize the images for subsequent analysis. The number of SVs at a particular synapse was quantified within four 50 nm shells (in nm: 50-100, 100-150, 150-200, 200-250) surrounding the T-bar (Hallermann et al., 2010b). Micrographs were acquired with 85,000-fold magnification.

#### Mouse RB terminal

The preparation of retinae from 8-10 weeks old cpx3^KO^ and wt mice has been described in (Mortensen et al., 2016) (Supplemental Experimental Procedures). Briefly, retinae were fixed in a cocktail of 4% paraformaldehyde and 2.5% glutaraldehyde in 0.1M phosphate buffer, embedded in low-melt agarose, and sectioned into 100 μm slices using a vibratome. Retina slices were processed by high-pressure freezing (Leica HPM100) and automated freeze substitution (Leica EM AFS2), and embedded in plastic for ultramicrotomy. Electron micrographs were acquired on a transmission electron microscope (Zeiss LEO 912-Omega, 80 kV) with 25,000-fold magnification. SVs surrounding ribbons of wt and *cpx3^KO^* RB synapses were counted within 200 nm concentric shells (in nm: 0-200, 200-400, 400-600, 600-800, 800-1000) centred at the ribbon base. The cytoplasmic area covered by each shell was also quantified to take into account cell morphology at presynaptic sites. In Figure 6, image contrast was manually adjusted to ease visualization.

### Synthesis and coupling of Brp peptides and pull-down assays

A 30-amino acid peptide (CTSVVPFPGGGGGQ<u>QQQQQDAGPAGFLKSFF</u>) including Brp^C-tip^ (underlined) and a scrambled control peptide(CTSVVPFPGGGGG<u>PFQQSLGFKAQQAQDQFG</u>) were synthesized by standard solid phase peptide synthesis using Fmoc chemistry. Peptides were immobilized to iodoacetyl agarose via an artificial cysteine residue added to their N-terminus.

*Drosophila* heads (from ~ 30 000 *w^1118^* animals) were homogenized in solubilization buffer [150 mM NaCl, 10 mM Hepes-NaOH (pH 7,4), 1 mM EGTA, 2 mM MgCl2, 1% NP-40, 1 mM DTT, Protease inhibitor cocktail] using an Ultra-Turrax (IKA), solubilized for 20 min (4°C) and centrifuged at 16,000x g for 30 min at 4°C. Next, the supernatant was collected into a fresh tube and proteins solubilized for another 10 min at 4°C before 15 min ultracentrifugation at 346,000× g. The resulting supernatant was partitioned, mixed with 100 μl functionalized beads and incubated for 3h at 4°C. Subsequently, beads were washed five times using solubilization buffer, resuspended in 1x Lämmli sample buffer and boiled for 5 min.

### Western blots

Fly heads were collected in standard radioimmunoprecipitation assay buffer [RIPA buffer; 150 mM NaCl, 1% Triton X-100, 0.5% sodium deoxycholate, 0.1% SDS, 50 mM Tris [pH 8.0)] supplemented with protease inhibitor cocktail (1:1000; Sigma-Aldrich) and immediately frozen in liquid nitrogen. Next, heads were homogenized using a pipette tip, supplemented with SDS-based protein buffer (Li-cor) and 2-mercaptoethanol (Merck). After a brief vortexing step, samples were centrifuged for 5 min at 13,000 rpm (4°C), incubated for 10 min at 55°C, subjected to electrophoresis on a 4-12% SDS gel and blotted onto 0.2 μm nitrocellulose membrane (Amersham™Protran™). The membrane was blocked for 1h using Odyssey Blocking buffer (Li-cor) diluted 1:8 with 1xPBS. Blots were probed with primary antisera at the indicated concentration for 1h at RT: rabbit-α-Cpx (1:10,000, (Huntwork and Littleton, 2007) and mouse-α-tubulin/*β* (1:1,000; Developmental Studies Hybridoma Bank e7, RRID: AB_528499). After rinsing twice and 3×10 min washing steps, membranes were incubated with IRDye 680RD goat-α-rabbit (RRID: AB_10956166) and 800CW goat-α-mouse (1:16,000, Li-cor, RRID: AB_621842) for 1h at RT, rinsed twice and washed 3×10 min. Proteins extracted from *w^1118^* fly head homogenate via pull-down experiments were separated by SDS-PAGE (18% Gel), immunoblotted using rabbit-α-Cpx (1:10,000 Huntwork and Littleton, 2007, RRID: AB_2568068) and goat-α-rabbit Alexa680 secondary antibody (1:5,000, Life). Western blots were imaged with an OdysseyFc 2800 (Li-cor).

### Electrophysiology

Two electrode voltage clamp (TEVC) recordings (Axoclamp 900A amplifier, Molecular Devices) were made from muscle 6, segments A2 and A3 of late third instar male *Drosophila* larvae essentially as previously described (Ljaschenko et al., 2013). All measurements were obtained at RT in HL-3 with the following composition (in mM): NaCl 70, KCl 5, MgCl_2_ 20, NHCO_3_ 10, trehalose 5, sucrose 115, HEPES 5, CaCl_2_ 1.5, 1 or 0.6 (as indicated), pH adjusted to 7.2. The intracellular electrodes had resistances of 10-20 MΩ, filled with 3M KCl. For analysis, only cells with an initial membrane potential of at least −50 mV and a membrane resistance of ≥ 4 MΩ were included. During recordings, cells were clamped at a holding potential of - 80 mV (minis) or −60 mV (eEPSCs). To evoke synaptic currents, nerves were stimulated via a suction electrode (diameter ~15 μm) with 300μs pulses, typically at 10 V (Grass S88 stimulator and isolation unit SIU5, Astro-Med). Signals were low-pass filtered at 10 kHz and analyzed in Clampfit 10.2 (Molecular Devices). Paired-pulse recordings were done with inter-stimulus intervals of (in ms): 10, 30, 100, 300, 1000. Between recordings, cells were given 10 seconds rest. For analysis, ten traces per interval were averaged. The amplitude of the second response in 10 ms inter-pulse recordings was measured from the peak to the point of interception with the extrapolated first eEPSC. To measure τ decay, the decaying phase of evoked currents was fitted with a mono-exponential function from 60% of the peak amplitude to the end of the event. In order to estimate RRV pool sizes, a train of 100 pulses was applied at 60 Hz. Linear fits from 0.3-0.5 s or 0.9-1.6 s were applied to cumulatively plotted eEPSCs and back-extrapolated (Hallermann et al., 2010a; Weyhersmuller et al., 2011).

### Larval locomotion

Briefly, wandering third instar larvae were positioned in a petri dish (9 cm in diameter) filled with 1 % agarose. The crawling paths of each genotype were recorded for 2 min using a digital camera and path lengths of individual animals were subsequently measured in ImageJ.

### Statistics

Data were analyzed with Prism 5.0 (GraphPad) and Sigma Plot 12.5 (Software Inc.). Group means were compared by a two-tailed Student’s t-test, unless the assumption of normality of the sample distribution was violated. In this case group means were compared by a non-parametric rank sum test. Data are reported as mean ± SEM, n indicates the sample number and P denotes the level of significance (*P ≤ 0.05, **P ≤ 0.01, and ***P ≤ 0.001). Statistics of all experiments are summarized in Tables S1, S2, S4-S11.

## SUPPLEMENTAL INFORMATION

**Figure S1, related to Figure 1.**
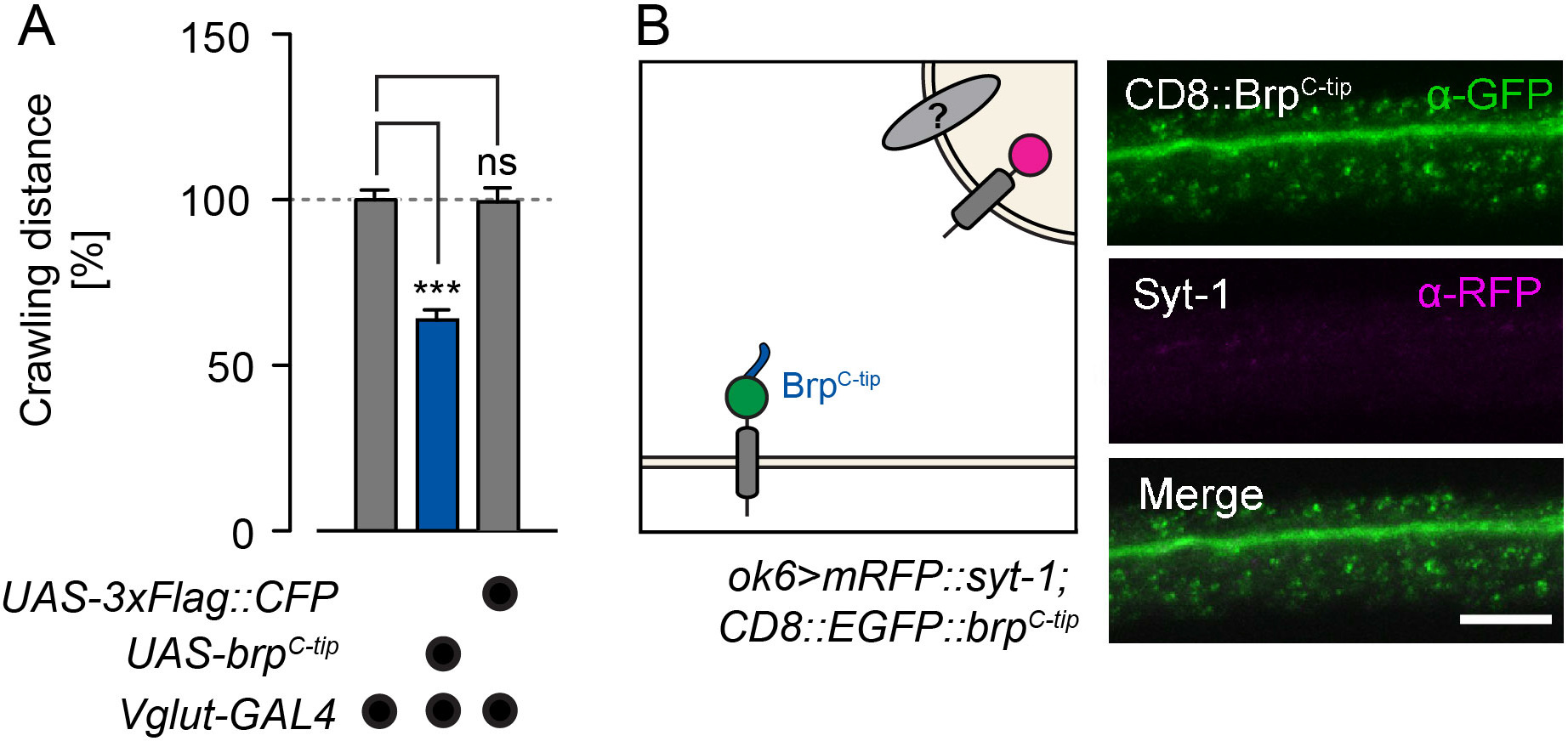
CFP is not responsible for SV tethering by cytoplasmicBrp^C-tip^. **(A)** Quantification of larval crawling distances shows comparable values for *Vglut-GAL4>UAS-3xFlag::CFP* and undriven *UAS-3xFlag::CFP* controls. Thus, the 3xFlag::CFP fusion is not responsible for the effect of Brp^C-tip^ (*3xFlag::CFP::brp^C-tip^)*. Scores are normalized to *Vglut-GAL4/+*. **(B)** Schematic Illustration (left) and confocal images (right) of motor axons expressing CD8::EGFP::Brp^C-tip^ (*ok6-GAL4>UAS-CD8::EGFP::brp^C-tip^*). Membrane-attached Brp^C-tip^ does not capture SVs at ectopic sites. Data presented as mean ± SEM, *** P ≤ 0.001. Scale bar 5 μm.

**Figure S2, related to Figure 2.**
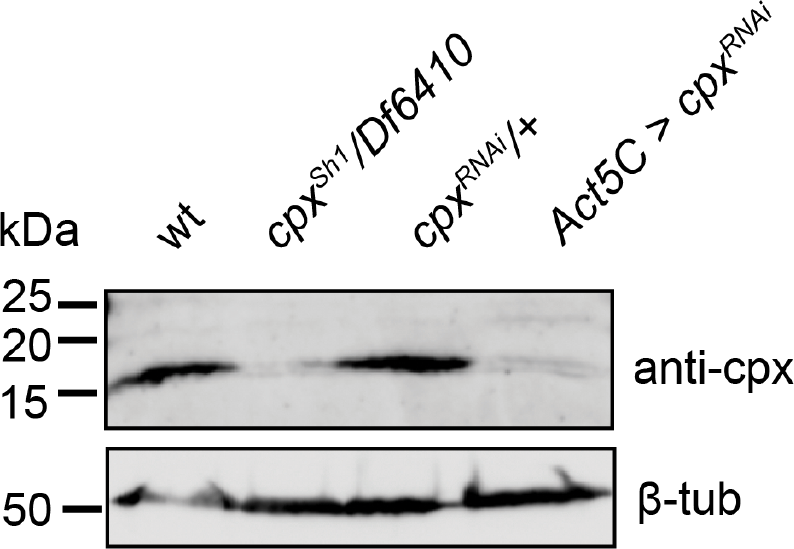
Cpx expression levels. The *cpx^RNAi^* strain employed in the screen for Brp interactors reduces *cpx* expression. Western blot analysis using a polyclonal antibody against Cpx (Huntwork and Littleton, 2007) shows absence and reduction of Cpx protein levels (~20 kDa band) in *cpx^Sh1^/Df6410* and *Actin-5C-GAL4>UAS-cpx^RNAi^* animals, respectively. β-tubulin served as loading control.

**Figure S3, related to Figure 2.**
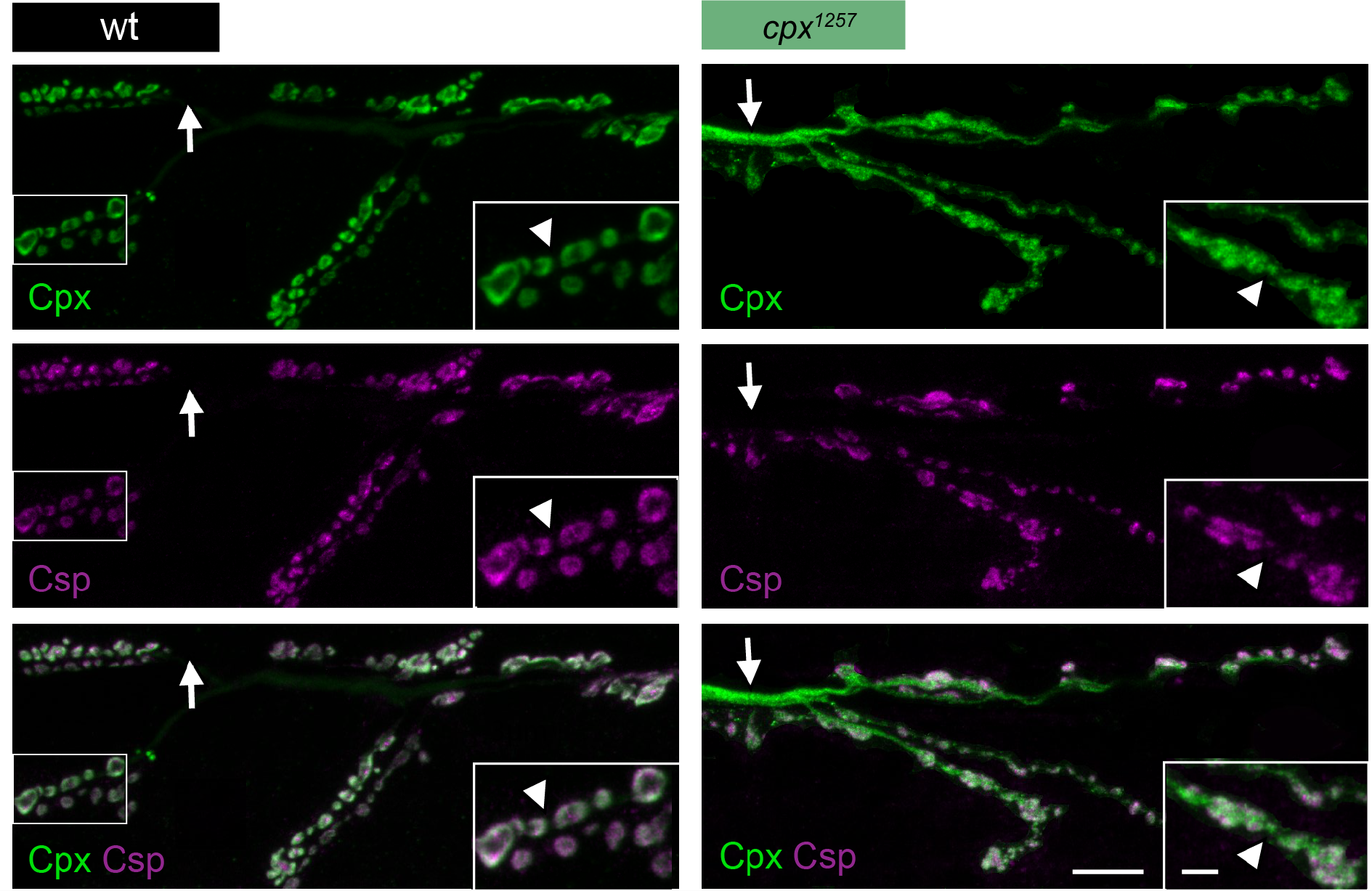
Disrupted farnesylation alters the distribution of Cpx at the NMJ. Co-staining against Cpx (green) and CSP (magenta) shows disturbed co-localization of Cpx with SVs at *cpx^1257^* NMJs (laser power was increased for Cpx imaging in mutants). Here, Cpx is distributed throughout motor axons (arrows) and inter-bouton sections (arrow heads) and no longer displays the characteristic enrichment in the bouton cortex. Scale bar 10 μm, inset 3 μm.

**Figure S4 related to Figure 2.**
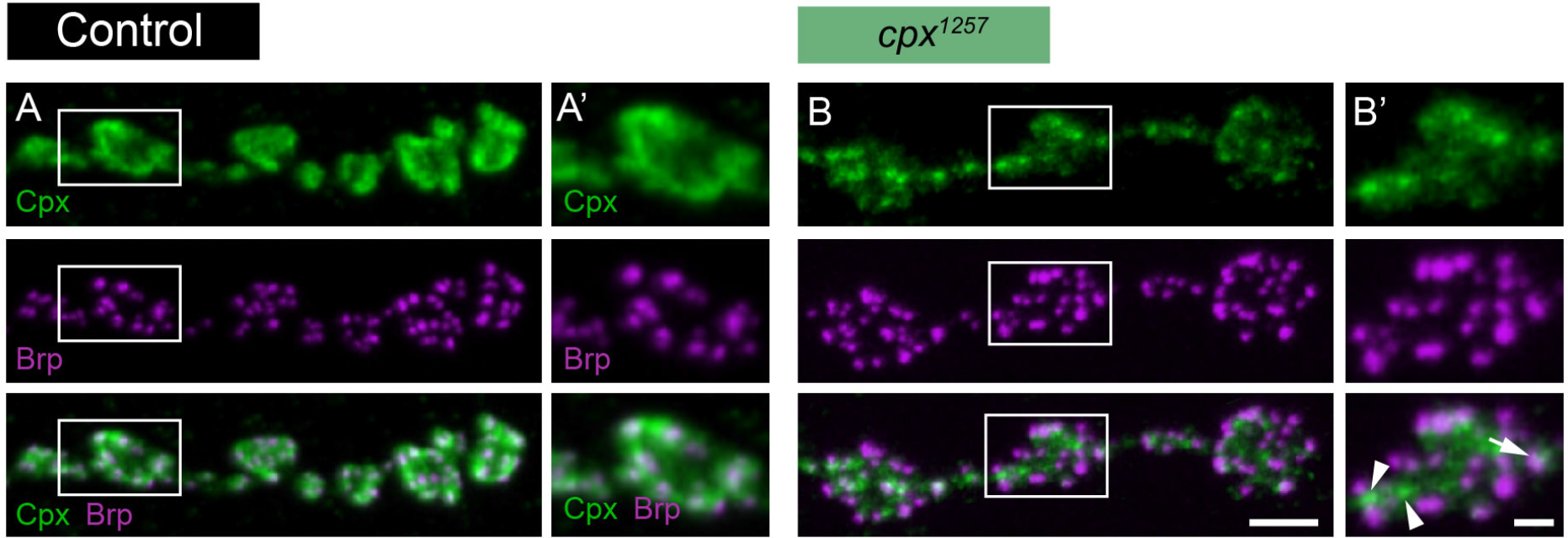
Cpx forms clusters at *cpx^1257^* mutant NMJs. Co-staining against Cpx (green) and Brp (magenta) at **(A, A’)** wt and **(B. B’)** *cpx^1257^* NMJs (laser power was increased for Cpx imaging in mutants). In the mutant, Cpx forms clusters, which are not visible in the wt and only partially overlap with active zones (arrow, co-localization with Brp; arrowheads, no overlap). Scale bars (A, B) 3 μm, (A’, B’) 1 μm.

**Figure S5, related to Figure 5.**
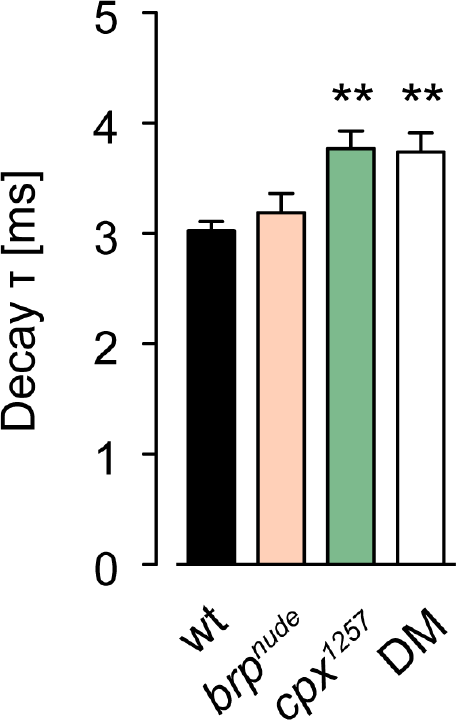
Decay kinetics of eEPSCs. The decay time constant (τ) of eEPSCs is unaltered in *brp^nude^* single mutants and prolonged in *cpx^1257^* animals and double mutants. This indicates that pronounced receptor desensitization does not contribute to the mutants’ electrophysiological phenotypes. Data presented as mean ± SEM, **P ≤ 0.01.

**Figure S6, related to Figure 5.**
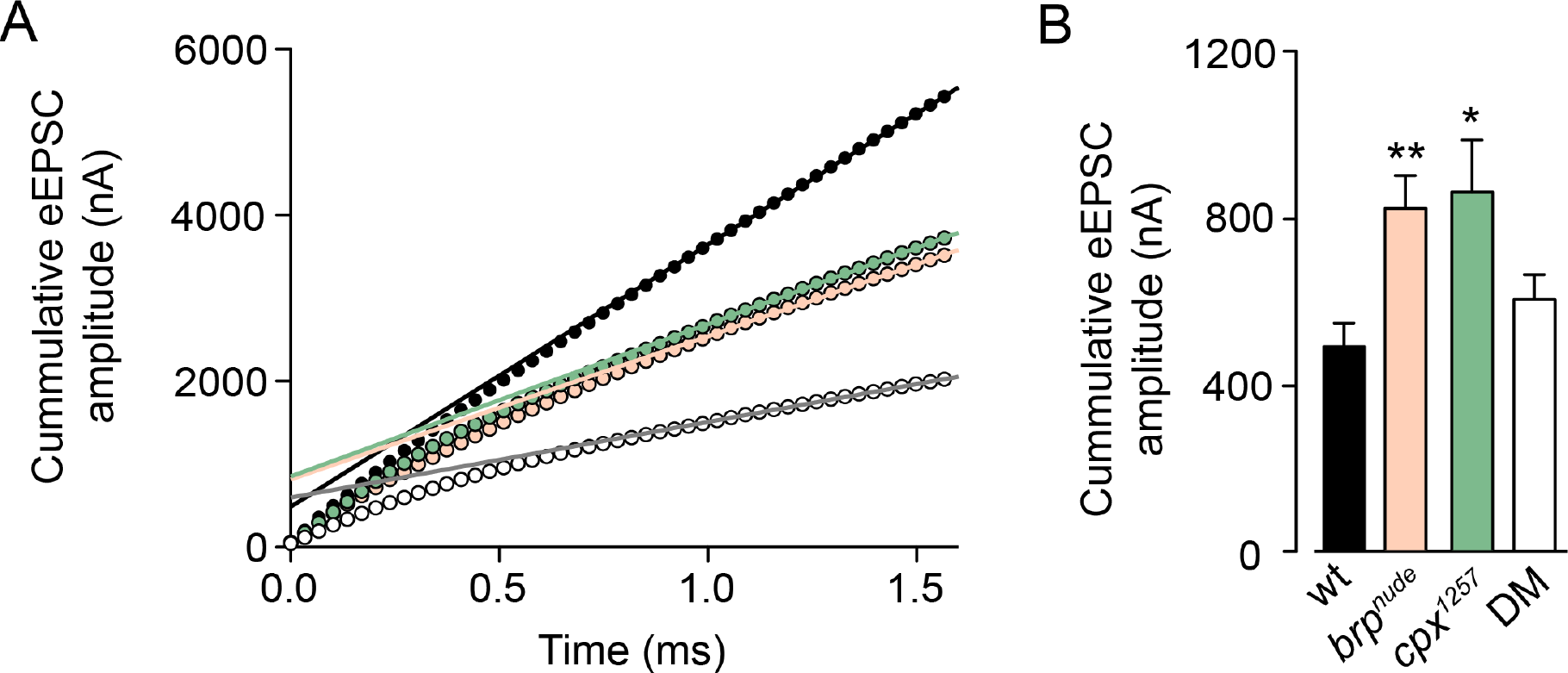
Estimating RRV pool sizes by later placement of the linear fit in the stimulus train. **(A)** Back-extrapolation of linear fits (0.9-1.6 s) to cumulatively plotted average eEPSC amplitudes at 60 Hz stimulation. **(B)** Alternative estimate (cf. **Figure 5I**) of RRV pool sizes in the different genotypes. Data are presented as mean ± S.E.M, * P ≤ 0.05, ** P ≤ 0.01.

**Figure S7, related to Figure 5.**
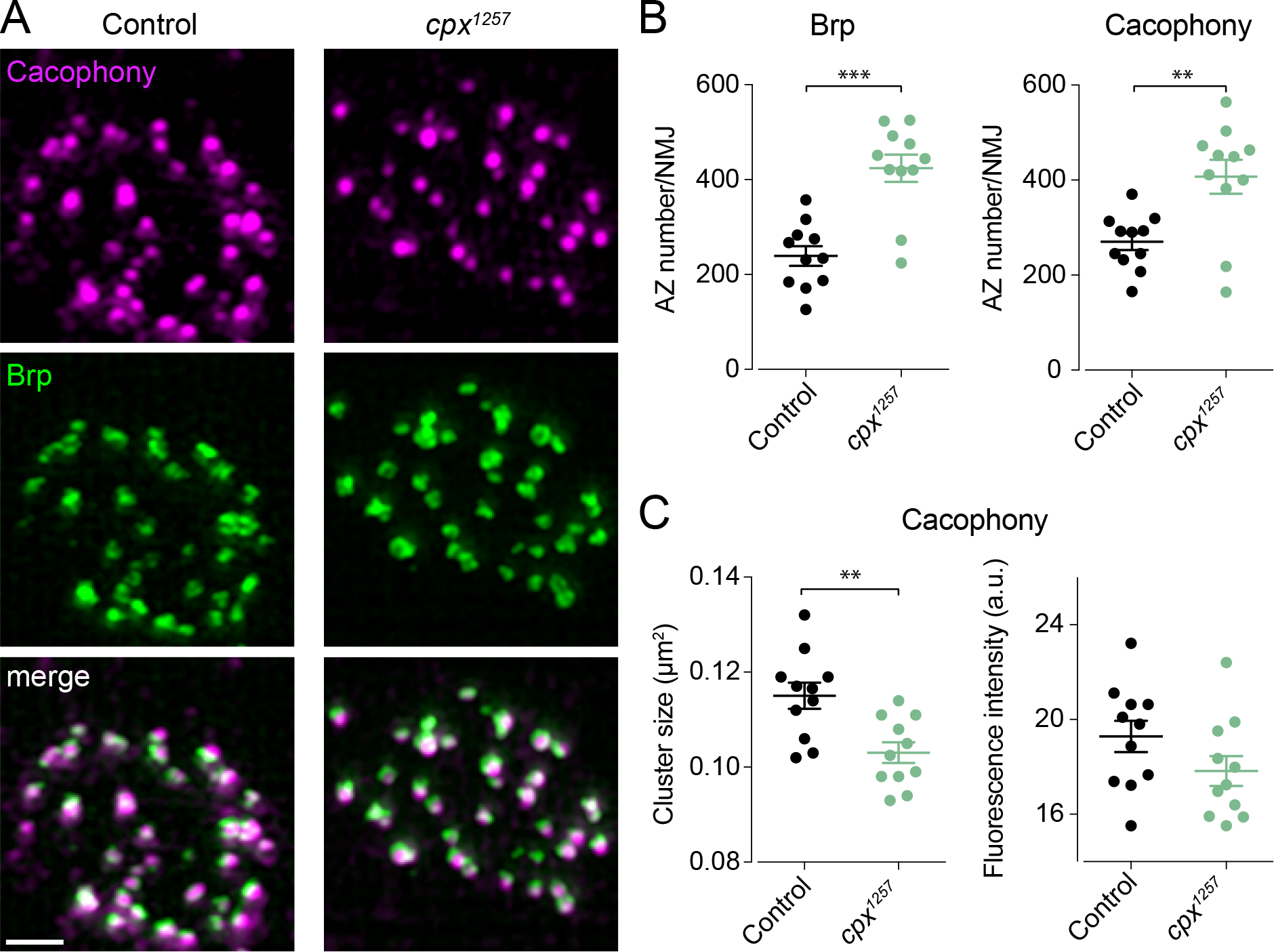
Ca^2+^-channel clusters at *cpx^1257^* active zones. **(A)** A GFP-tagged version of Cacophony, an α1 subunit of voltage-gated Ca^2+^-channels (Kawasaki et al., 2004; Kittel et al., 2006), was expressed in larval motoneurons (*ok6-GAL4>UAS-cac^GFP^*) and visualized by SIM. Examples show stainings against GFP (magenta) and Brp (mAb nc82, green). **(B)** Quantification of imaging data from muscle 4 shows an increased number of Cacophony clusters and Brp-positive active zones in *cpx^1257^* mutants. Whereas their signal intensity is unaltered, the average size of Cacophony clusters is significantly reduced in *cpx^1257^* mutants. Data are presented as mean ± S.E.M, ** P ≤ 0.01, *** P ≤ 0.001. Scale bar 1 μm.

**Figure S8, related to Figure 6.**
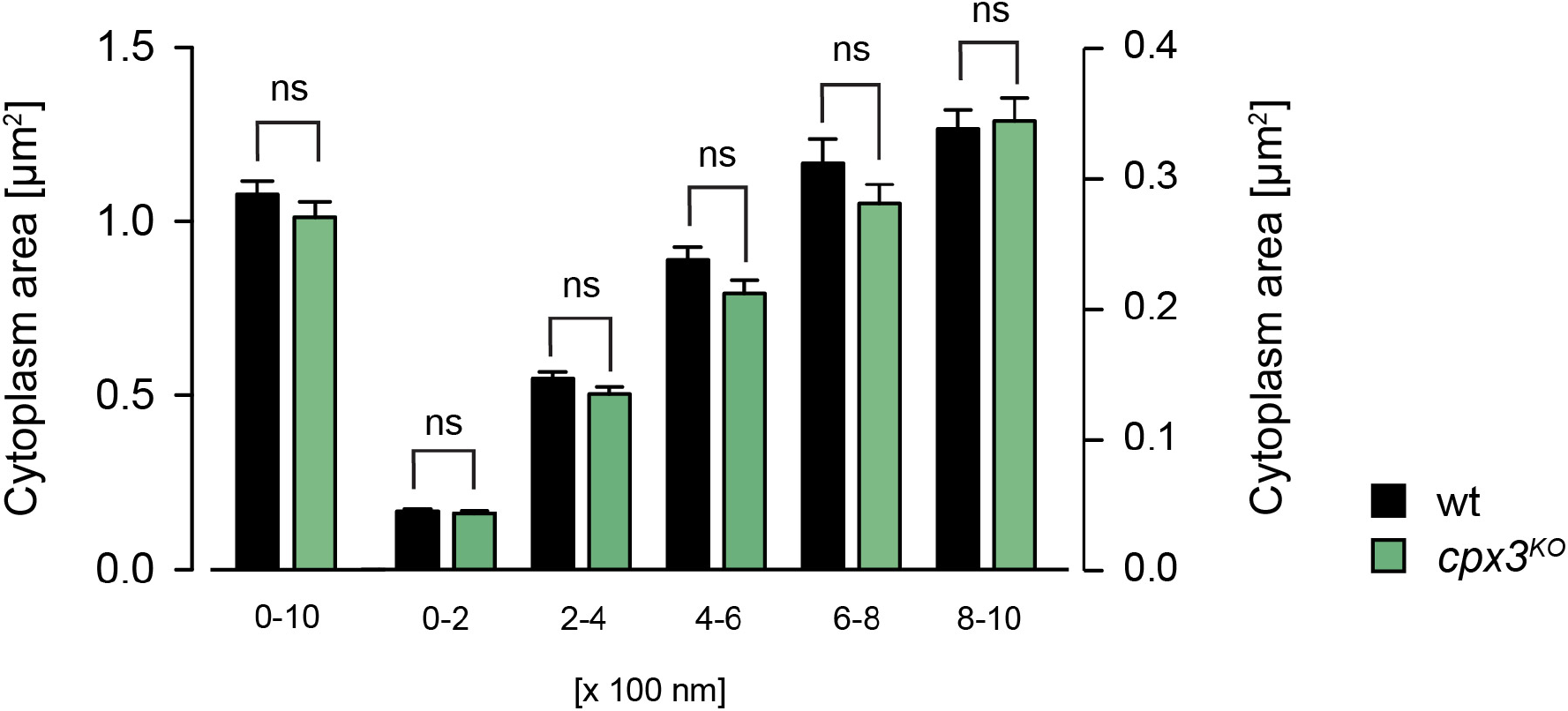
Ribbons of *cpx3^KO^* RB terminals are surrounded by a normal cytoplasmic space. Analysis of electron micrographs of wt (black) and *cpx3^KO^* (green) RB terminals. Quantification of the cytoplasmic area surrounding synaptic ribbons within five 200 nm thick concentric shells (right axis) and in sum (left axis, 0-1000 nm). Data are presented as mean ± S.E.M.

**Figure S9.**
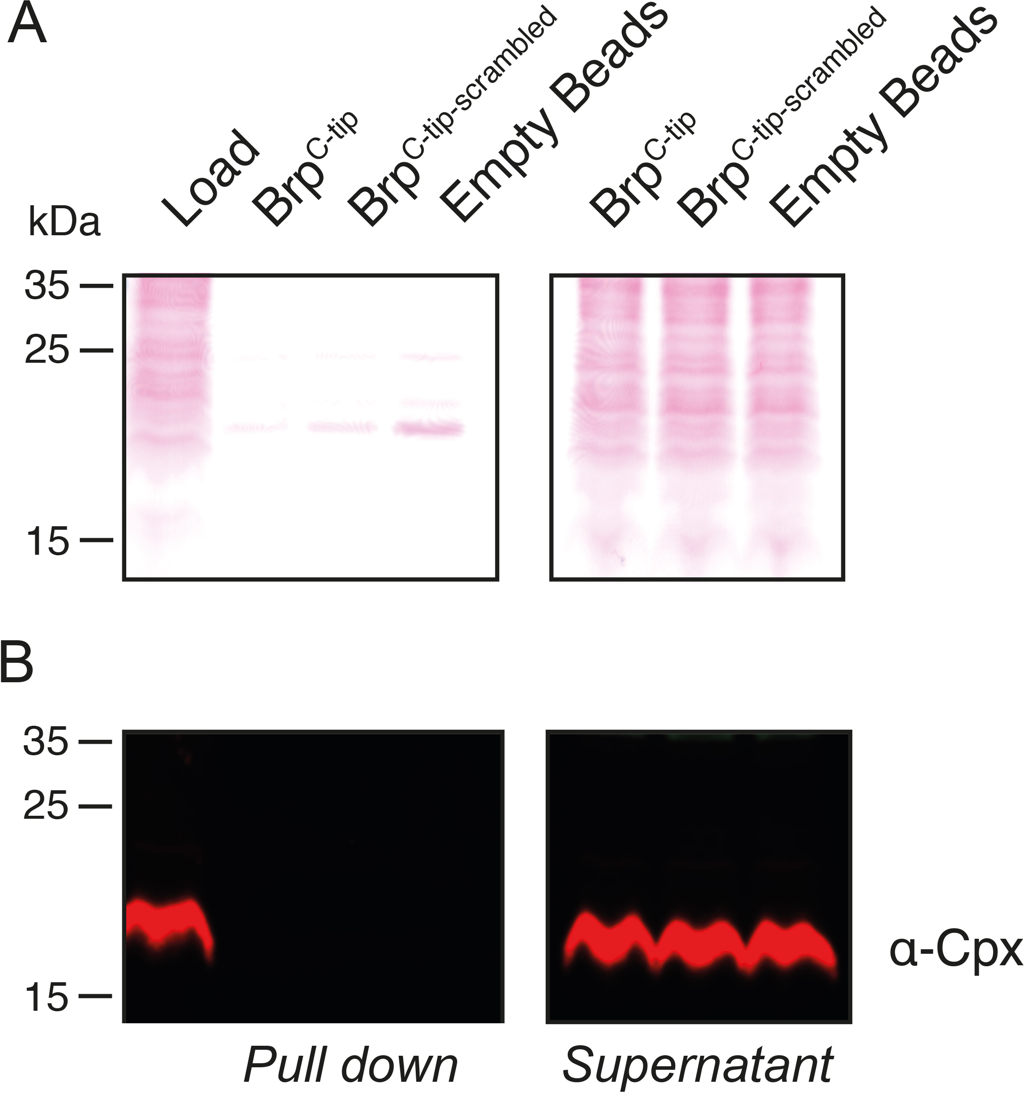
The interaction between Cpx and Brp^C-tip^ escapes biochemical capture. A synthetic peptide including Brp^C-tip^ was immobilized to generate an affinity matrix for capturing Cpx. A Brp^C-tip^ variant with randomly scrambled amino acid sequence and empty agarose beads served as controls. **(A)** Ponceau-S membrane staining. Left panel: Proteins enriched by immobilized Brp^C-tip^. Right panel: Unbound protein. **(B)** Western blot analysis. Cpx was detected in the load (~20 kDa band) suggesting sufficient protein extraction but was not found in the eluate from the affinity matrix containing Brp^C-tip^.

**Table S1.**
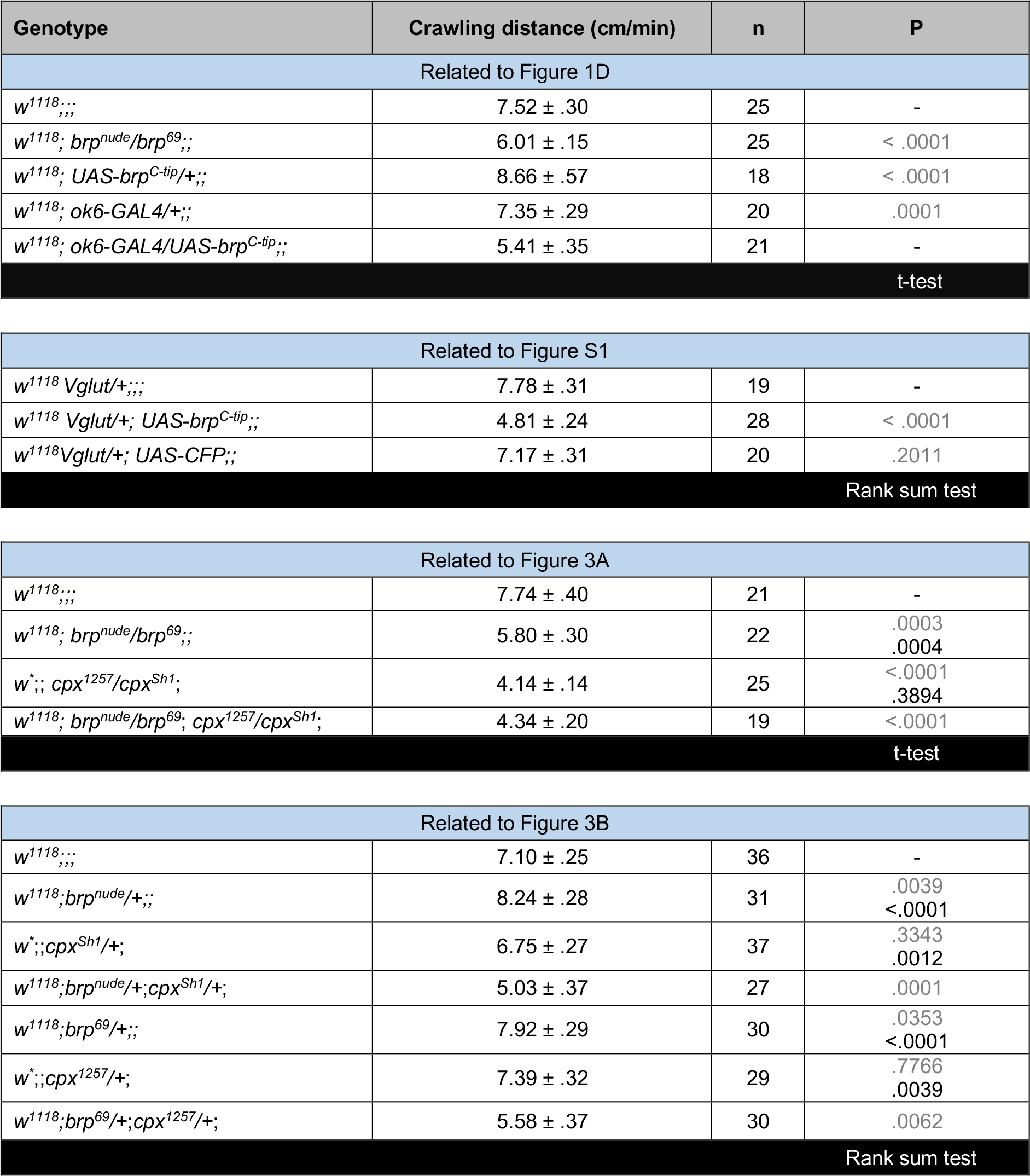
Crawling distance. Values represent the mean ± SEM, n denotes number of larvae. P values in grey and black indicate comparisons with genetic controls and between experimental groups, respectively (as indicated in the figures).

| Genotype | Crawling distance (cm/min) | n | P |
| --- | --- | --- | --- |
| Related to Figure 1D |  |  |  |
| *w<sup>1118</sup>...* | 7.52 ± .30 | 25 | - |
| *w<sup>1118</sup>; brp<sup>nude</sup>/brp<sup>69</sup>;* | 6.01 ± .15 | 25 | < .0001 |
| *w<sup>1118</sup>; UAS-brp<sup>C-tip</sup>/+;* | 8.66 ± .57 | 18 | < .0001 |
| *w<sup>1118</sup>; ok6-GAL4/+;* | 7.35 ± .29 | 20 | .0001 |
| *w<sup>1118</sup>; ok6-GAL4/UAS-brp<sup>C-tip</sup>;* | 5.41 ± .35 | 21 | - |
| t-test |  |  |  |
| Related to Figure S1 |  |  |  |
| *w<sup>1118</sup> Vglut/+;* | 7.78 ± .31 | 19 | - |
| *w<sup>1118</sup> Vglut/+; UAS-brp<sup>C-tip</sup>;* | 4.81 ± .24 | 28 | < .0001 |
| *w<sup>1118</sup> Vglut/+; UAS-CFP;* | 7.17 ± .31 | 20 | .2011 |
| Rank sum test |  |  |  |
| Related to Figure 3A |  |  |  |
| *w<sup>1118</sup>...* | 7.74 ± .40 | 21 | - |
| *w<sup>1118</sup>; brp<sup>nude</sup>/brp<sup>69</sup>;* | 5.80 ± .30 | 22 | .0003<br>.0004 |
| *w<sup>\*</sup>; cpx<sup>1257</sup>/cpx<sup>Sh1</sup>;* | 4.14 ± .14 | 25 | <.0001<br>.3894 |
| *w<sup>1118</sup>; brp<sup>nude</sup>/brp<sup>69</sup>; cpx<sup>1257</sup>/cpx<sup>Sh1</sup>;* | 4.34 ± .20 | 19 | <.0001 |
| t-test |  |  |  |
| Related to Figure 3B |  |  |  |
| *w<sup>1118</sup>...* | 7.10 ± .25 | 36 | - |
| *w<sup>1118</sup>; brp<sup>nude</sup>/+;* | 8.24 ± .28 | 31 | .0039<br><.0001 |
| *w<sup>\*</sup>; cpx<sup>Sh1</sup>/+;* | 6.75 ± .27 | 37 | .3343<br>.0012 |
| *w<sup>1118</sup>; brp<sup>nude</sup>/+; cpx<sup>Sh1</sup>/+;* | 5.03 ± .37 | 27 | .0001 |
| *w<sup>1118</sup>; brp<sup>69</sup>/+;* | 7.92 ± .29 | 30 | .0353<br><.0001 |
| *w<sup>\*</sup>; cpx<sup>1257</sup>/+;* | 7.39 ± .32 | 29 | .7766<br>.0039 |
| *w<sup>1118</sup>; brp<sup>69</sup>/+; cpx<sup>1257</sup>/+;* | 5.58 ± .37 | 30 | .0062 |
| Rank sum test |  |  |  |

**Table S2.**
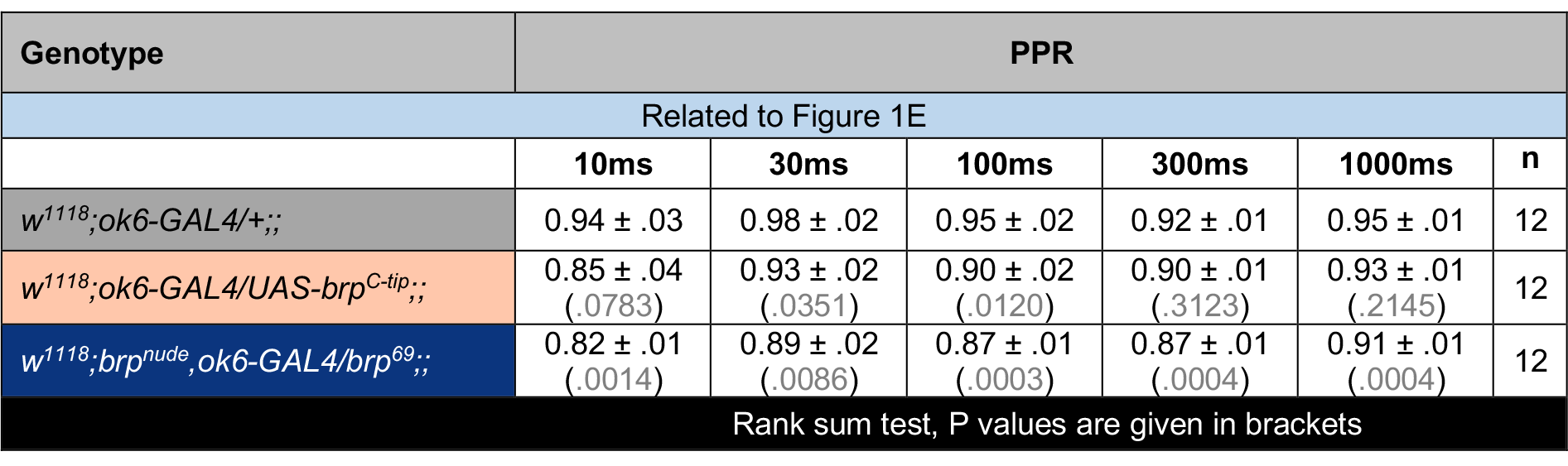
Quantification of paired-pulse ratios (PPR) following Brp^C-tip^ expression. Data are presented as mean ± SEM, n denotes number of NMJs. P values indicate comparisons with the control (*w^1118^;ok6-GAL4/+;;)*.

**Table S3.** Candidates for the genetic screen. List of 27 different genes tested in the RNAi-based screen. The gene products are involved in different steps of the SV cycle.

| Functional involvement | Symbol | Protein name | Off-targets | Chromosome | VDRC stock |
| --- | --- | --- | --- | --- | --- |
| Related to Figure 2A |  |  |  |  |  |
| **Priming/**<br>**regulation** | *dunc-13* | dunc-13 | 0 | 2 | v33606 |
|  | *tomosyn* | tomosyn | 0 | 3 | v43629 |
|  | *cpx* | complexin | 0 | 3 | v21477 |
| **Fusion/Ca<sup>2+</sup>-sensing** | *dCirl* | Ca<sup>2+</sup>-independent receptor of latrotoxin | 110 | 2 | v29969 |
|  | *dCAPS* | CAPS | 0 | 2 | v25291 |
|  | *csp* | Cysteine string protein | 1 | 2 | v103201 |
|  | *syt-12* | Synaptotagmin-12 | 0 | 2 | V47506 |
|  | *syt-7* | Synaptotagmin-7 | 0 | 3 | v24988 |
| **Regulators,**<br>**effectors & Rabs** | *rim-1* | Rab3-interacting molecule | 0 | 2 | v39384 |
|  | *rab3-GAP* | rab3 GTPase binding protein | 0 | 3 | v27824 |
|  | *gdi* | GDP dissociation inhibitor | 1 | 3 | v26537 |
|  | *rph* | Rabphilin | 15 | 2 | v52438 |
|  | *rac-1* |  | 1 | 1 | v49247 |
|  | *rab5* | Rab3-binding protein 5 | 3 | 2 | v34094 |
| **SV phospho-proteins** | *sng-1* | Synaptogyrin | 0 | 3 | v8784 |
|  | *syn* | Synapsin | 0 | 2 | v110606 |
| **Endocytosis** | *AnnB9* | AnnexinB9 | 2 | 2 | v27493 |
|  | *AnnB10* | AnnexinB10 | 0 | 2 | v36107 |
|  | *twf* | Twinfilin | 0 | 2 | v25817 |
| **Others** | *dysb* | Dysbindin | 0 | 2 | v34354 |
|  | *atg-1* | Autophagy-specific gene 1 | 0 | 2 | v16133 |
|  | *rsk* | Ribosomal S6 kinase | 14 | 3 | v5702 |
|  | *sap47* | Synapse-associated proteine 47 kDa | 0 | 2 | v35445 |
|  | *dsyd-1* | Sunday driver-1 | 0 | 2 | v35346 |
|  | *dvglut* | Vesicular glutamat transporter | 0 | 3 | v2574 |
|  | *vti1a* | Vesicle transport through interaction with t-SNAREs homolog 1A | 0 | 3 | v45726 |
|  | *Mctp* | Multiple-C2-domain TM proteins | 0 | 3 | v10061 |

**Table S4.**
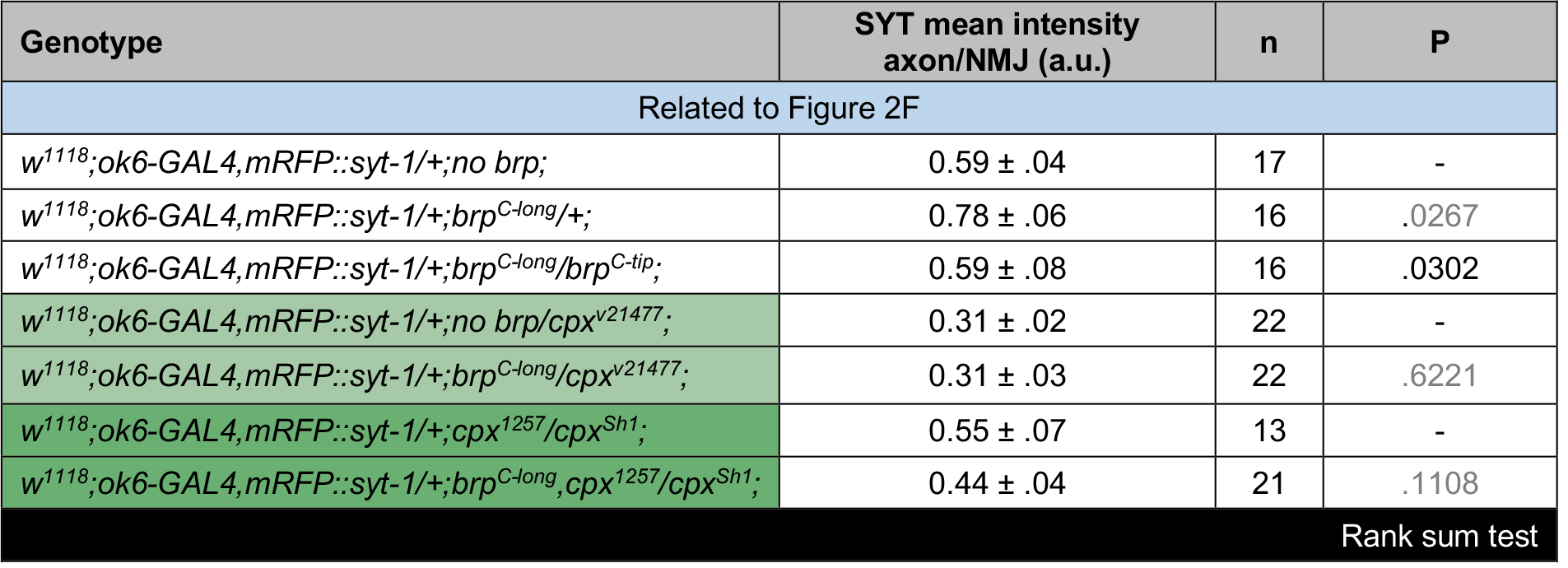
Imaging basis to screen for Brp interactors. SV distributions (axon/NMJ ratio) in wt (white), *cpx^v12477(RNAi)^* (light green) and *cpx^1257^* (dark green) genetic backgrounds. Values represent the mean ± SEM, n denotes number of larvae. P values in grey and black indicate comparisons with genetic controls and between experimental groups, respectively.

**Table S5.** Quantification of SV numbers at NMJ T-bars. Values represent the mean ± SEM, n denotes number of T-bars. P values in grey and black indicate comparisons with wt and the double mutant, respectively.

| Genotype | Number of SVs/T-bar |  |  |  |  |
| --- | --- | --- | --- | --- | --- |
| Related to Figure 4B |  |  |  |  |  |
|  | 50-100nm | 100-150nm | 150-200nm | 200-250nm | n |
| *w<sup>1118...</sup><sub>,,,</sub>* | 6.0 ± .19 | 8.14 ± .28 | 8.93 ± .43 | 11.76 ± .54 | 58 |
| *w<sup>1118</sup>;*brp*<sup>nude</sup>/*brp*<sup>69...</sup><sub>,,</sub>* | 3.40 ± .22<br>(*<.0001*)<br>(.8933) | 5.36 ± .26<br>(*<.0001*)<br>(.8407) | 6.92 ± .29<br>(.0004)<br>(.1143) | 10.62 ± .48<br>(.0700)<br>(.5105) | 42 |
| *w<sup>\*</sup>;*cpx*<sup>1257</sup>/*cpx*<sup>Sh1</sup>;* | 4.52 ± .22<br>(*<.0001*)<br>(.0034) | 5.80 ± .20<br>(*<.0001*)<br>(.1553) | 6.91 ± .27<br>(.0001)<br>(.1052) | 10.64 ± .43<br>(.0732)<br>(.4081) | 65 |
| *w<sup>1118</sup>;*brp*<sup>nude</sup>/*brp*<sup>69</sup>;*cpx*<sup>1257</sup>/*cpx*<sup>Sh1</sup>;* | 3.50 ± .22<br>(*<.0001*) | 5.24 ± .27<br>(*<.0001*) | 6.24 ± .37<br>(*<.0001*) | 10.11 ± .60<br>(.0325) | 38 |
| Rank sum test, P values are given in brackets |  |  |  |  |  |

**Table S6.** *d*STORM analysis of the CAZ ultrastructure. Data are represented as mean ± SEM, n denotes number of NMJs. P values indicate comparisons with wt.

| Genotype | CAZ area ($\mu\text{m}^2$) | P | Localizations/CAZ | P | n |
| --- | --- | --- | --- | --- | --- |
| Related to Figure 4C |  |  |  |  |  |
| *w<sup>1118</sup>...* | 0.09 ± .003 | - | 556.39 ± 48.81 | - | 19 |
| *w<sup>1118</sup>;brp<sup>nude</sup>/brp<sup>69</sup>;;* | 0.07 ± .002 | .0002 | 552.89 ± 42.19 | 1.00 | 21 |
| *w<sup>\*</sup>;;cp<sup>x</sup><sup>1257</sup>/cp<sup>x</sup><sup>Sh1</sup>;* | 0.08 ± .003 | .0507 | 534.57 ± 53.19 | .8673 | 18 |
| *w<sup>1118</sup>;brp<sup>nude</sup>/brp<sup>69</sup>;cp<sup>x</sup><sup>1257</sup>/cp<sup>x</sup><sup>Sh1</sup>;* | 0.08 ± .002 | .0016 | 578.37 ± 51.19 | .8331 | 20 |
| t-test Rank sum test |  |  |  |  |  |

**Table S7.** Electrophysiological analysis of mutant NMJs. Data are presented as mean ± SEM, n denotes number of NMJs. P values in grey and black indicate comparisons with wt and the double mutant, respectively.

| Genotype | eEPSC amplitude (-nA) | P | n | Mini frequency (Hz) | P | n |
| --- | --- | --- | --- | --- | --- | --- |
| Related to Figure 5E |  |  | Related to Figure 5B |  |  |  |
| *w<sup>1118...</sup>* | 44.80 ± 3.02 | - | 10 | 1.11 ± .18 | - | 15 |
| *w<sup>1118</sup>;brp<sup>nude</sup>/brp<sup>69</sup>;* | 50.50 ± 5.05 | .3800<br>.5924 | 13 | 0.83 ± .05 | .4450<br>< .0001 | 14 |
| *w<sup>\*</sup>;cpx<sup>1257</sup>/cpx<sup>Sh1</sup>;* | 52.77 ± 2.28 | .0431<br>.1858 | 13 | 62.60 ± 5.91 | < .0001<br>< .0001 | 14 |
| *w<sup>1118</sup>;brp<sup>nude</sup>/brp<sup>69</sup>;cpx<sup>1257</sup>/cpx<sup>Sh1</sup>;* | 47.09 ± 3.57 | .6374 | 12 | 23.19 ± 2.52 | < .0001 | 14 |
| t-test |  |  | Rank sum test |  |  |  |

| Genotype | PPR (1 mM Ca<sup>2+</sup>) |  |  |  |  |  |
| --- | --- | --- | --- | --- | --- | --- |
| Related to Figure 5G |  |  |  |  |  |  |
|  | 10ms | 30ms | 100ms | 300ms | 1000ms | n |
| *w<sup>1118</sup>...* | 1.50 ± .10 | 1.51 ± .11 | 1.19 ± .04 | 1.05 ± .02 | 0.99 ± .01 | 10 |
| *w<sup>1118</sup>;brp<sup>nude</sup>/brp<sup>69</sup>...* | 1.08 ± .07<br>(.0039) | 1.21 ± .04<br>(.0101) | 0.97 ± .02<br>(.0200) | 1.01 ± .02<br>(.1138) | 0.98 ± .01<br>(.4757) | 13 |
| *w<sup>\*</sup>;;cpx<sup>1257</sup>/cpx<sup>Sh1</sup>;* | 1.03 ± .03<br>(.0009) | 1.05 ± .03<br>(.0002) | 0.98 ± .02<br>(.0006) | 0.94 ± .02<br>(.0006) | 0.96 ± .01<br>(.0586) | 13 |
| *w<sup>1118</sup>;brp<sup>nude</sup>/brp<sup>69</sup>;cpx<sup>1257</sup>/cpx<sup>Sh1</sup>;* | 0.80 ± .06<br>(.0002) | 0.90 ± .04<br>(.0001) | 0.85 ± .02<br>(.0001) | 0.87 ± .02<br>(.0001) | 0.91 ± .01<br>(.0011) | 12 |
| Rank sum test, P values are given in brackets |  |  |  |  |  |  |

| Genotype | PPR (0.6 mM Ca<sup>2+</sup>) |  |  |  |  |  |
| --- | --- | --- | --- | --- | --- | --- |
| Related to Figure 5G |  |  |  |  |  |  |
|  | 10ms | 30ms | 100ms | 300ms | 1000ms | n |
| *w<sup>1118...</sup><sub>,,,</sub>* | 1.81 ± .09 | 1.62 ± .08 | 1.30 ± .05 | 1.12 ± .05 | 0.10 ± .03 | 14 |
| *w<sup>1118</sup>;brp<sup>nude</sup>/brp<sup>69</sup>...* | 1.58 ± .09<br>(.0769) | 1.45 ± .08<br>(.1354) | 1.24 ± .07<br>(.1238) | 1.18 ± .03<br>(.0565) | 1.00 ± .02<br>(.9817) | 14 |
| *w<sup>\*</sup>;cpx<sup>1257</sup>/cpx<sup>Sh1</sup>;* | 1.44 ± .09<br>(.0123) | 1.27 ± .09<br>(.0019) | 1.07 ± .03<br>(.0026) | 1.07 ± .05<br>(.1354) | 0.98 ± .02<br>(.8362) | 14 |
| *w<sup>1118</sup>;brp<sup>nude</sup>/brp<sup>69</sup>;cpx<sup>1257</sup>/cpx<sup>Sh1</sup>;* | 1.25 ± .07<br>(.0004) | 1.15 ± .05<br>(.0002) | 1.00 ± .02<br>(.0005) | 0.94 ± .02<br>(.0013) | 0.95 ± .02<br>(.4520) | 13 |
| Rank sum test, P values are given in brackets |  |  |  |  |  |  |

| Genotype | eEPSC decay $\tau$ (ms) | P | n |
| --- | --- | --- | --- |
| Related to Figure S5 |  |  |  |
| *w<sup>1118...</sup>* | 3.03 ± 0.08 | - | 10 |
| *w<sup>1118</sup>;brp<sup>nude</sup>/brp<sup>69</sup>;* | 3.19 ± 0.17 | .4462 | 13 |
| *w<sup>\*</sup>;cpx<sup>1257</sup>/cpx<sup>Sh1</sup>;* | 3.77 ± 0.16 | .0011 | 13 |
| *w<sup>1118</sup>;brp<sup>nude</sup>/brp<sup>69</sup>;cpx<sup>1257</sup>/cpx<sup>Sh1</sup>;* | 3.74 ± 0.17 | .0022 | 12 |
| t-test |  |  |  |

**Table S8.** Quantification of AZ numbers. Data are represented as mean ± SEM, n denotes number of NMJs. P values in grey and black indicate comparisons with wt and the double mutant, respectively.

| Genotype | AZ number | P | n |
| --- | --- | --- | --- |
| Related to Figure 5C |  |  |  |
| *w<sup>1118...</sup>* | 414.06 ± 26.59 | - | 16 |
| *w<sup>1118</sup>;brp<sup>nude</sup>/brp<sup>69</sup>;;* | 352.50 ± 26.42 | .1683<br>.8968 | 8 |
| *w<sup>\*</sup>;;cpx<sup>1257</sup>/cpx<sup>Sh1</sup>;* | 523.78 ± 32.16 | .0138<br>.0133 | 9 |
| *w<sup>1118</sup>;brp<sup>nude</sup>/brp<sup>69</sup>;cpx<sup>1257</sup>/cpx<sup>Sh1</sup>;* | 369.10 ± 27.64 | .2357 | 10 |
| Rank sum test |  |  |  |

**Table S9.** Back-extrapolation of cumulatively plotted eEPSC amplitudes. Estimates of average RRV pool sizes using linear fits to 0.3-0.5 s (upper table) or 0.9-1.6 s (lower table) of the stimulus train. Data are presented as mean ± SEM, n denotes number of NMJs. P values in grey and black indicate comparisons with wt and the double mutant, respectively.

| Genotype | Cumulative eEPSC amplitude (nA) | P | n |
| --- | --- | --- | --- |
| Related to Figure 5I |  |  |  |
| *w<sup>1118...</sup>* | 205.77 ± 39.10 | - | 9 |
| *w<sup>1118</sup>;brp<sup>nude</sup>/brp<sup>69</sup>;;* | 242.66 ± 32.58 | .4771<br>.2442 | 13 |
| *w<sup>\*</sup>;;cpx<sup>1257</sup>/cpx<sup>Sh1</sup>;* | 335.59 ± 37.10 | .0292<br>.0032 | 13 |
| *w<sup>1118</sup>;brp<sup>nude</sup>/brp<sup>69</sup>;cpx<sup>1257</sup>/cpx<sup>Sh1</sup>;* | 197.72 ± 16.80 | .8380 | 12 |
| t-test |  |  |  |
| Related to Figure S6 |  |  |  |
| *w<sup>1118...</sup>* | 489.79 ± 56.14 | - | 9 |
| *w<sup>1118</sup>;brp<sup>nude</sup>/brp<sup>69</sup>;;* | 815.39 ± 78.36 | .0033<br>.0240 | 13 |
| *w<sup>\*</sup>;;cpx<sup>1257</sup>/cpx<sup>Sh1</sup>;* | 854.49 ± 124.22 | .0194<br>.1654 | 13 |
| *w<sup>1118</sup>;brp<sup>nude</sup>/brp<sup>69</sup>;cpx<sup>1257</sup>/cpx<sup>Sh1</sup>;* | 598.56 ± 58.82 | .1886 | 12 |
| Rank sum test |  |  |  |

**Table S10.**
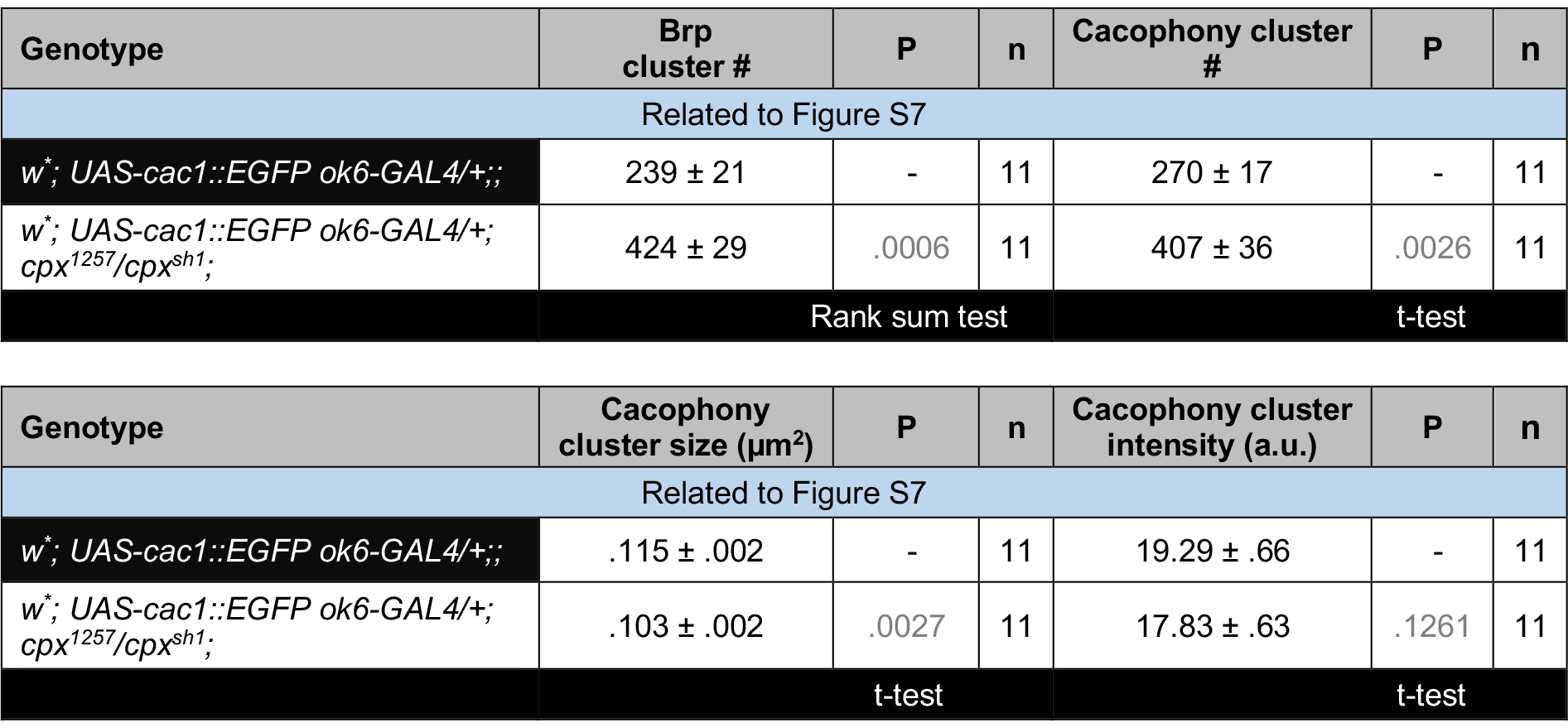
Quantification of Ca^2+^-channel clusters at *cpx^1257^* active zones. Data are presented as mean ± SEM, n denotes number of NMJs.

**Table S11.** Analysis of synaptic ribbons at mouse RB terminals. Data are represented as mean ± SEM, n denotes number of ribbons. P values indicate comparisons with wt.

| Genotype | SV numbers/200 nm shell |  |  |  |  |  |
| --- | --- | --- | --- | --- | --- | --- |
| Related to Figure 6C |  |  |  |  |  |  |
|  | 0-200 nm | 200-400 nm | 400-600 nm | 600-800 nm | 800-1000 nm | n |
| wt | 4.76 ± .34 | 17.00 ± 1.01 | 26.83 ± 1.83 | 29.91 ± 2.10 | 34.48 ± 2.36 | 46 |
| *cpx3<sup>ko</sup>* | 3.00 ± .22<br>(.0001) | 13.44 ± 0.74<br>(.0042) | 22.42 ± 1.62<br>(.1069) | 27.46 ± 1.77<br>(.3987) | 33.60 ± 2.12<br>(.8589) | 48 |
| Rank sum test, P values are given in brackets |  |  |  |  |  |  |

| Genotype | Cytoplasm area (μm<sup>2</sup>)/200 nm shell |  |  |  |  |  |
| --- | --- | --- | --- | --- | --- | --- |
| Related to Figure S8 |  |  |  |  |  |  |
|  | 0-200 nm | 200-400 nm | 400-600 nm | 600-800 nm | 800-1000 nm | n |
| wt | 0.05 ± .002 | 0.15 ± .005 | 0.24 ± .010 | 0.31 ± .018 | 0.34 ± .015 | 46 |
| *cpx3<sup>ko</sup>* | 0.04 ± .002<br>(.2515) | 0.13 ± .005<br>(.0581) | 0.21 ± .010<br>(.1433) | 0.28 ± .015<br>(.3311) | 0.34 ± .018<br>(.8264) | 48 |
| Rank sum test, P values are given in brackets |  |  |  |  |  |  |

## REFERENCES

Brand, A.H., and Perrimon, N. (1993). Targeted gene expression as a means of altering cell fates and generating dominant phenotypes. Development 118, 401–415.

Brose, N. (2008). For better or for worse: Complexins regulate SNARE function and vesicle fusion. Traffic 9, 1403–1413.

Buhl, L.K., Jorquera, R.A., Akbergenova, Y., Huntwork-Rodriguez, S., Volfson, D., and Littleton, J.T. (2013). Differential regulation of evoked and spontaneous neurotransmitter release by C-terminal modifications of complexin. Mol. Cell. Neurosci. 52, 161–172.

Cho, R.W., Song, Y., and Littleton, J.T. (2010). Comparative analysis of Drosophila and mammalian complexins as fusion clamps and facilitators of neurotransmitter release. Mol. Cell. Neurosci. 45, 389–397.

Choi, B.J., Imlach, W.L., Jiao, W., Wolfram, V., Wu, Y., Grbic, M., Cela, C., Baines, R.A., Nitabach, M.N., and McCabe, B.D. (2014). Miniature neurotransmission regulates Drosophila synaptic structural maturation. Neuron 82, 618–634.

Crawford, D.C., and Kavalali, E.T. (2015). Molecular underpinnings of synaptic vesicle pool heterogeneity. Traffic 16, 338–364.

Daniels, R.W. (2004). Increased expression of the Drosophila vesicular glutamate transporter leads to excess glutamate release and a compensatory decrease in quantal content. J. Neurosci. 24, 10466–10474.

Daniels, R.W., Gelfand, M. V., Collins, C.A., and DiAntonio, A. (2008). Visualizing glutamatergic cell bodies and synapses in Drosophila larval and adult CNS. J. Comp. Neurol. 508, 131–152.

Denker, a., Krohnert, K., Buckers, J., Neher, E., and Rizzoli, S.O. (2011). The reserve pool of synaptic vesicles acts as a buffer for proteins involved in synaptic vesicle recycling. Proc. Natl. Acad. Sci. 108, 17183–17188.

DiAntonio, A., Petersen, S.A., Heckmann, M., and Goodman, C.S. (1999). Glutamate receptor expression regulates quantal size and quantal content at the Drosophila neuromuscular junction. J. Neurosci. 19, 3023–3032.

Dietzl, G., Chen, D., Schnorrer, F., Su, K.C., Barinova, Y., Fellner, M., Gasser, B., Kinsey, K., Oppel, S., Scheiblauer, S., et al. (2007). A genome-wide transgenic RNAi library for conditional gene inactivation in Drosophila. Nature 448, 151–156.

Ehmann, N., van de Linde, S., Alon, A., Ljaschenko, D., Keung, X.Z., Holm, T., Rings, A., DiAntonio, A., Hallermann, S., Ashery, U., et al. (2014). Quantitative super-resolution imaging of Bruchpilot distinguishes active zone states. Nat. Commun. 5, 4650.

Fernández-Busnadiego, R., Asano, S., Oprisoreanu, A.M., Sakata, E., Doengi, M., Kochovski, Z., Zürner, M., Stein, V., Schoch, S., Baumeister, W., et al. (2013). Cryo-electron tomography reveals a critical role of RIM1 a in synaptic vesicle tethering. J. Cell Biol. 201, 725–740.

Fouquet, W., Owald, D., Wichmann, C., Mertel, S., Depner, H., Dyba, M., Hallermann, S., Kittel, R.J., Eimer, S., and Sigrist, S.J. (2009). Maturation of active zone assembly by Drosophila Bruchpilot. J. Cell Biol. 186, 129–145.

Frank, T., Rutherford, M.A., Strenzke, N., Neef, A., Pangr??i??, T., Khimich, D., Fetjova, A., Gundelfinger, E.D., Liberman, M.C., Harke, B., et al. (2010). Bassoon and the synaptic ribbon organize Ca^2+^ channels and vesicles to add release sites and promote refilling. Neuron 68, 724–738.

Gong, J., Lai, Y., Li, X., Wang, M., Leitz, J., Hu, Y., Zhang, Y., Choi, U.B., Cipriano, D., Pfuetzner, R.A., et al. (2016). C-terminal domain of mammalian complexin-1 localizes to highly curved membranes. Proc. Natl. Acad. Sci. U. S. A. 349, 475–481.

Groth, A.C., Fish, M., Nusse, R., and Calos, M.P. (2004). Construction of Transgenic Drosophila by Using the Site-Specific Integrase from Phage φC31. Genetics 166, 1775–1782.

Gustafsson, M.G.L., Shao, L., Carlton, P.M., Wang, C.J.R., Golubovskaya, I.N., Cande, W.Z., Agard, D.A., and Sedat, J.W. (2008). Three-dimensional resolution doubling in wide-field fluorescence microscopy by structured illumination. Biophys. J. 94, 4957–4970.

Hallermann, S., and Silver, R.A. (2013). Sustaining rapid vesicular release at active zones: potential roles for vesicle tethering. Trends Neurosci. 36, 185–194.

Hallermann, S., Heckmann, M., and Kittel, R.J. (2010a). Mechanisms of short-term plasticity at neuromuscular active zones of Drosophila. HFSP J. 4, 72–84.

Hallermann, S., Kittel, R.J., Wichmann, C., Weyhersmuller, A., Fouquet, W., Mertel, S., Owald, D., Eimer, S., Depner, H., Schwarzel, M., et al. (2010b). Naked dense bodies provoke depression. J. Neurosci. 30, 14340–14345.

Hallermann, S., Fejtova, A., Schmidt, H., Weyhersmüller, A., Silver, R.A., Gundelfinger, E.D., and Eilers, J. (2010c). Bassoon speeds vesicle reloading at a central excitatory synapse. Neuron 68, 710–723.

Heckmann, M., and Dudel, J. (1997). Desensitization and resensitization kinetics of glutamate receptor channels from Drosophila larval muscle. Biophys. J. 72, 2160–2169.

Heilemann, M., Van De Linde, S., Schüttpelz, M., Kasper, R., Seefeldt, B., Mukherjee, A., Tinnefeld, P., and Sauer, M. (2008). Subdiffraction-resolution fluorescence imaging with conventional fluorescent probes. Angew. Chemie - Int. Ed. 47, 6172–6176.

Hobson, R.J., Liu, Q., Watanabe, S., and Jorgensen, E.M. (2011). Complexin maintains vesicles in the primed state in C. elegans. Curr. Biol. 21, 106–113.

Hosoi, N., Holt, M., and Sakaba, T. (2009). Calcium dependence of exo- and endocytotic coupling at a glutamatergic synapse. Neuron 63, 216–229.

Huntwork, S., and Littleton, J.T. (2007). A complexin fusion clamp regulates spontaneous neurotransmitter release and synaptic growth. Nat. Neurosci. 10, 1235–1237.

Ito, K., Awano, W., Suzuki, K., Hiromi, Y., and Yamamoto, D. (1997). The Drosophila mushroom body is a quadruple structure of clonal units each of which contains a virtually identical set of neurones and glial cells. Development 124, 761–771.

Iyer, J., Wahlmark, C.J., Kuser-Ahnert, G.A., and Kawasaki, F. (2013). Molecular mechanisms of COMPLEXIN fusion clamp function in synaptic exocytosis revealed in a new Drosophila mutant. Mol. Cell. Neurosci. 56, 244–254.

Jockusch, W.J., Speidel, D., Sigler, A., Sørensen, J.B., Varoqueaux, F., Rhee, J.S., and Brose, N. (2007). CAPS-1 and CAPS-2 are essential synaptic vesicle priming proteins. Cell 131, 796–808.

Kaeser, P.S., and Regehr, W.G. (2014). Molecular mechanisms for synchronous, asynchronous, and spontaneous neurotransmitter release. Annu. Rev. Physiol. 76, 333–363.

Kawasaki, F., Hazen, M., and Ordway, R.W. (2000). Fast synaptic fatigue in shibire mutants reveals a rapid requirement for dynamin in synaptic vesicle membrane trafficking. Nat. Neurosci. 3, 859–860.

Kawasaki, F., Zou, B., Xu, X., and Ordway, R.W. (2004). Active zone localization of presynaptic calcium channels encoded by the cacophony locus of Drosophila. J. Neurosci. 24, 282–285.

Kittel, R.J., Wichmann, C., Rasse, T.M., Fouquet, W., Schmidt, M., Schmid, A., Wagh, D.A., Pawlu,C., Kellner, R.R., Willig, K.I., et al. (2006). Bruchpilot promotes active zone assembly, Ca^2+^ channel clustering, and vesicle release. Science 312, 1051–1054.

van de Linde, S., Löschberger, A., Klein, T., Heidbreder, M., Wolter, S., Heilemann, M., and Sauer, M. (2011). Direct stochastic optical reconstruction microscopy with standard fluorescent probes. Nat. Protoc. 6, 991–1009.

Ljaschenko, D., Ehmann, N., and Kittel, R.J. (2013). Hebbian plasticity guides maturation of glutamate receptor fields in vivo. Cell Rep. 3, 1407–1413.

Markstein, M., Pitsouli, C., Villalta, C., Celniker, S.E., and Perrimon, N. (2008). Exploiting position effects and the gypsy retrovirus insulator to engineer precisely expressed transgenes. Nat. Genet. 40, 476–483.

Martin, J.A., Hu, Z., Fenz, K.M., Fernández, J., and Dittman, J.S. (2011). Complexin has opposite effects on two modes of synaptic vesicle fusion. Curr. Biol. 21, 97–105.

Melom, J.E., Akbergenova, Y., Gavornik, J.P., and Littleton, J.T. (2013). Spontaneous and evoked release are independently regulated at individual active zones. J. Neurosci. 33, 17253–17263.

Midorikawa, M., and Sakaba, T. (2015). Imaging exocytosis of single synaptic vesicles at a fast CNS presynaptic terminal. Neuron 88, 492–498.

Miskiewicz, K., Jose, L.E., Yeshaw, W.M., Valadas, J.S., Swerts, J., Munck, S., Feiguin, F., Dermaut, B., and Verstreken, P. (2014). HDAC6 is a Bruchpilot deacetylase that facilitates neurotransmitter release. Cell Rep. 8, 94–102.

Mortensen, L.S., Park, S.J.H., Ke, J. Bin, Cooper, B.H., Zhang, L., Imig, C., Löwel, S., Reim, K., Brose, N., Demb, J.B., et al. (2016). Complexin 3 increases the fidelity of signaling in a retinal circuit by regulating exocytosis at ribbon synapses. Cell Rep. 15, 2239–2250.

Neher, E. (2010a). Complexin: does it deserve its name? Neuron 68, 803–806.

Neher, E. (2010b). What is rate-limiting during sustained synaptic activity: vesicle supply or the availability of release sites. Front. Synaptic Neurosci. 2, 1–6.

Neher, E. (2015). Merits and limitations of vesicle pool models in view of heterogeneous populations of synaptic vesicles. Neuron 87, 1131–1142.

Paul, M.M., Pauli, M., Ehmann, N., Hallermann, S., Sauer, M., Kittel, R.J., and Heckmann, M. (2015). Bruchpilot and Synaptotagmin collaborate to drive rapid glutamate release and active zone differentiation. Front. Cell. Neurosci. 9, 1–12.

Reim, K. (2017). Complexins. In Reference Module in Neuroscience and Biobehavioral Psychology (Elsevier Inc.).

Reim, K., Wegmeyer, H., Brandstätter, J.H., Xue, M., Rosenmund, C., Dresbach, T., Hofmann, K., and Brose, N. (2005). Structurally and functionally unique complexins at retinal ribbon synapses. J. Cell Biol. 169, 669–680.

Reim, K., Regus-Leidig, H., Ammermüller, J., El-Kordi, A., Radyushkin, K., Ehrenreich, H., Brandstätter, J.H., and Brose, N. (2009). Aberrant function and structure of retinal ribbon synapses in the absence of complexin 3 and complexin 4. J. Cell Sci. 122, 1352–1361.

Reynolds, E.S. (1963). The use of lead citrate stain at high pH in electron microscopy. J. Cell Biol. 17, 208.

Rizzoli, S.O., and Betz, W.J. (2004). The structural organization of the readily releasable pool of synaptic vesicles. Science 303, 2037–2039.

Sabeva, N., Cho, R.W., Vasin, A., Gonzalez, A., Littleton, J.T., and Bykhovskaia, M. (2017). Complexin mutants reveal partial segregation between recycling pathways that drive evoked and spontaneous neurotransmission. J. Neurosci. 37, 383–396.

Sanyal, S. (2009). Genomic mapping and expression patterns of C380, OK6 and D42 enhancer trap lines in the larval nervous system of Drosophila. Gene Expr. Patterns 9, 371–380.

Schäfer, P., Van De Linde, S., Lehmann, J., Sauer, M., and Doose, S. (2013). Methylene blue- and thiol-based oxygen depletion for super-resolution imaging. Anal. Chem. 85, 3393–3400.

Schmid, A., Hallermann, S., Kittel, R.J., Khorramshahi, O., Frölich, A.M.J., Quentin, C., Rasse, T.M., Mertel, S., Heckmann, M., and Sigrist, S.J. (2008). Activity-dependent site-specific changes of glutamate receptor composition in vivo. Nat. Neurosci. 11, 659–666.

Schneggenburger, R., Meyer, A.C., and Neher, E. (1999). Released fraction and total size of a pool of immediately available transmitter quanta at a calyx synapse. Neuron 23, 399–409.

Sharonov, A., and Hochstrasser, R.M. (2007). Single-molecule imaging of the association of the cell-penetrating peptide Pep-1 to model membranes. Biochemistry 46, 7963–7972.

Snellman, J., Mehta, B., Babai, N., Bartoletti, T.M., Akmentin, W., Francis, A., Matthews, G., Thoreson, W., and Zenisek, D. (2011). Acute destruction of the synaptic ribbon reveals a role for the ribbon in vesicle priming. Nat. Neurosci. 14, 1135–1141.

Stewart, B.A., Atwood, H.L., Renger, J.J., Wang, J., and Wu, C.F. (1994). Improved stability of Drosophila larval neuromuscular preparations in haemolymph-like physiological solutions. J. Comp. Physiol. A 175, 179–191.

Tokunaga, M., Imamoto, N., and Sakata-Sogawa, K. (2008). Highly inclined thin illumination enables clear single-molecule imaging in cells. Nat. Methods 5, 159–161.

Trimbuch, T., and Rosenmund, C. (2016). Should I stop or should I go? The role of complexin in neurotransmitter release. Nat. Rev. Neurosci. 17, 118–125.

Wagh, D.A., Rasse, T.M., Asan, E., Hofbauer, A., Schwenkert, I., Dürrbeck, H., Buchner, S., Dabauvalle, M.C., Schmidt, M., Qin, G., et al. (2006). Bruchpilot, a protein with homology to ELKS/CAST, is required for structural integrity and function of synaptic active zones in Drosophila. Neuron 49, 833–844.

Wang, S.S.H., Held, R.G., Wong, M.Y., Liu, C., Karakhanyan, A., and Kaeser, P.S. (2016). Fusion competent synaptic vesicles persist upon active zone disruption and loss of vesicle docking. Neuron 91, 777–791.

Weyhersmuller, A., Hallermann, S., Wagner, N., and Eilers, J. (2011). Rapid active zone remodeling during synaptic plasticity. J. Neurosci. 31, 6041–6052.

Wilhelm, B.G., Mandad, S., Truckenbrodt, S., Kröhnert, K., Schäfer, C., Rammner, B., Koo, S.J., Claßten, G. a, Krauss, M., Haucke, V., et al. (2014). Composition of isolated synaptic boutons reveals the amounts of vesicle trafficking proteins. Science 344, 1023–1028.

Wolter, S., Schüttpelz, M., Tscherepanow, M., Van De Linde, S., Heilemann, M., and Sauer, M. (2010). Real-time computation of subdiffraction-resolution fluorescence images. J. Microsc. 237, 12–22.

Wolter, S., Löschberger, A., Holm, T., Aufmkolk, S., Dabauvalle, M.C., Van De Linde, S., and Sauer, M. (2012). RapidSTORM: Accurate, fast open-source software for localization microscopy. Nat. Methods 9, 1040–1041.

Wragg, R.T., Snead, D., Dong, Y., Ramlall, T.F., Menon, I., Bai, J., Eliezer, D., and Dittman, J.S. (2013). Synaptic vesicles position Complexin to block spontaneous fusion. Neuron 77, 323–334.

Xue, M., Lin, Y.Q., Pan, H., Reim, K., Deng, H., Bellen, H.J., and Rosenmund, C. (2009). Tilting the balance between facilitatory and inhibitory functions of mammalian and Drosophila complexins crchestrates synaptic vesicle exocytosis. Neuron 64, 367–380.

Yook, K.J., Proulx, S.R., and Jorgensen, E.M. (2001). Rules of nonallelic noncomplementation at the synapse in Caenorhabditis elegans. Genetics 158, 209–220.

Zenisek, D., Steyer, J.A., and Almers, W. (2000). Transport, capture and exocytosis of single synaptic vesicles at active zones. Nature 406, 849–854.

Zhang, F.L., and Casey, P.J. (1996). Protein prenylation: molecular mechanisms and functional consequences. Annu. Rev. Biochem. 65, 241–269.

Zucker, R.S., and Regehr, W.G. (2002). Short-term synaptic plasticity. Annu. Rev. Physiol. 64, 355–405.

